# Adaptive replication origin activation alters chromatin dynamics and stability in cancer cells

**DOI:** 10.1101/2024.12.03.626323

**Authors:** Alejandro Pérez-Venteo, Fernando Unzueta, Sonia Feu, Pere Llinàs-Arias, Caroline Mauvezin, Neus Agell

**Author notes:** Both corresponding authors". Both authors had an equal contribution (Alejandro Pérez-Venteo, Fernando Unzueta).

## Abstract

Genome duplication, critical for cell survival and identity, requires precise origin activation to ensure accurate DNA replication and chromatin structure maintenance. In colorectal cancer cells, prolonged replication stress does not hinder cell proliferation, although the mechanisms driving cancer cell adaptability remain largely unclear. Here, we demonstrate that upon recovery, cancer cells are able to activate new replication origins in distinct domains, causing persistent changes in chromatin accessibility, nuclear morphology, and replication timing, ultimately promoting chromatin instability. Chromatin accessibility, particularly in promoter regions, increased following replication stress in a subset of cells, correlating with altered gene expression and nuclear expansion. Additionally, some genes with enhanced promoter accessibility displayed sustained expression changes, further suggesting a transcriptional shift linked to stress adaptation. Our findings reveal that colorectal cancer cells recover from severe replication stress through new origin activation, a mechanism that not only maintains cell proliferation under stress but may also accelerate tumour heterogeneity. This research underscores replication origin activation as a potential therapeutic target in combating cancer cell resilience.

**GRAPHICAL ABSTRACT:** 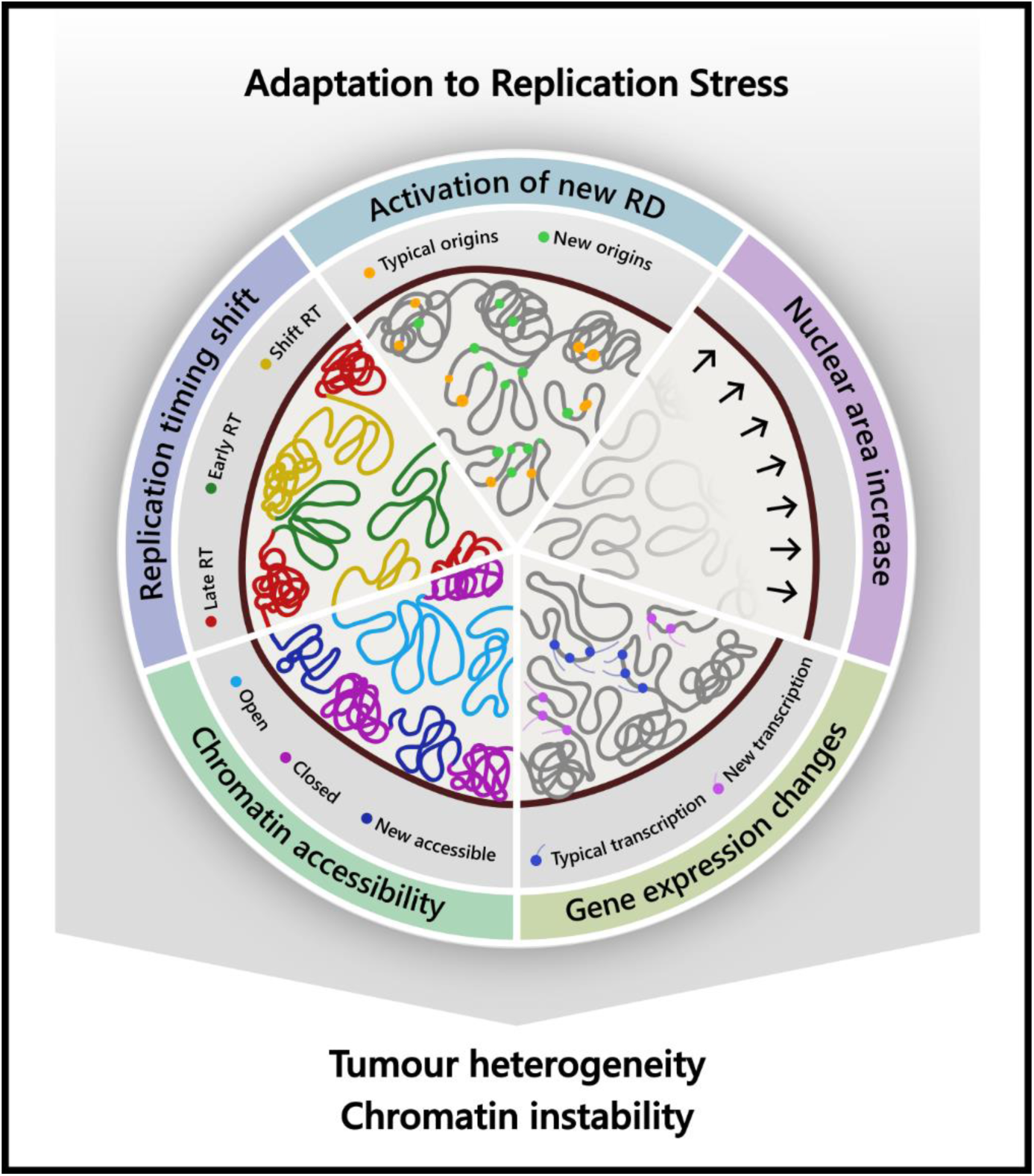

**MAIN HIGHLIGHTS:**

- Colorectal cancer cells under severe replication stress activate new replication origins upon recovery.
- The firing of new origins in distinct replication domains results in long-lasting changes in chromatin accessibility, especially in promoter regions, and is associated with alterations in nuclear morphology and replication timing.
- Increased chromatin accessibility in certain promoter regions correlates with prolonged gene expression changes.
- The capacity of CRC cells to recover from severe replication stress fosters chromatin instability and tumour heterogeneity.

## INTRODUCTION

Genome duplication is a crucial event of the cell cycle, not only for cell proliferation but also to maintain genome integrity and cell identity (Branzei & Foiani, 2010; Hanahan & Weinberg, 2011). During this process, DNA should be copied with very high fidelity and all DNA should be replicated once and only once. Additionally, chromatin features must be maintained and copied: new DNA helixes should have the same epigenetic marks as the parental ones and nuclear organization of chromatin in the nucleus must be preserved. Due to the large size of DNA in eukaryotic cells, its duplication is initiated at multiple replication origins located across all chromosomes. Interestingly, not all origins are activated simultaneously, instead they are triggered in groups according to a precise, yet not fully understood, replication timing program (Fragkos *et al*, 2015; Kang *et al*, 2018; Rivera-Mulia & Gilbert, 2016b; Aladjem & Redon, 2017). To prevent the reinitiation of replication at any origin, the processes of origin licensing and firing are strictly separated into two different cell cycle phases. Licensing occurs exclusively during the G1 phase, when the MCM-replicative helicase complex (MCM2-7) is loaded onto the DNA (Errico & Costanzo, 2010; Fragkos *et al*, 2015). Origin firing takes place in the S phase, where and when additional proteins are bound to origins, forming the Cdc45-MCM-GINS (CMG) complex. This complex activates the helicase, and consequently leads to the unwinding of DNA for replication. As previously mentioned above, origins are not fired simultaneously but in groups according to the replication timing program. Each activated origin generates two replication forks that move away in opposite directions, driven by separate replisomes containing DNA polymerases. During the G1 phase, more origins are licensed than are needed for replication, with most of them remaining unused as dormant origins. These dormant origins can serve as backups, ensuring complete genome replication if problems arise during DNA replication (Muñoz & Méndez, 2016; Técher *et al*, 2017; Ge *et al*, 2007). When newly synthetized DNA is immunolabeled and observed under a microscope in eukaryotic cells, hundreds of foci are visible, each containing multiple active replication forks (Chagin *et al*, 2016). These foci correspond to regions of DNA that share the same replication timing. Proper coordination of origin firing has been previously demonstrated to be essential for genome stability (Donley & Thayer, 2013; Neelsen *et al*, 2013; Alver *et al*, 2014). Additionally, in mammals, cell fate commitment is also associated with dynamic changes in replication timing within units known as replication domains (RDs), which range in size from approximately 400 to 800 kb (Hiratani *et al*, 2010, 2008; Ryba *et al*, 2010). These RD might correspond to the foci observed by immunofluorescence (Rivera-Mulia & Gilbert, 2016a; Wilson *et al*, 2016). The coordination of origin firing appears to play an important role in maintaining chromatin structure, although the underlying mechanisms and cause-consequence effect remain unclear.

Both extrinsic - such as DNA alkylating or crosslinking agents - and intrinsic factors - like dNTPs depletion, conflicts with the transcription machinery, or the formation of secondary DNA structures - can impair replisome progression and induce replication stress. It is now widely accepted that DNA replication stress is a major contributor of genomic instability and a key driver of tumorigenesis (Macheret & Halazonetis, 2015; Bester *et al*, 2011; Burrell *et al*, 2013a; Gaillard *et al*, 2015). To maintain genome stability, cells respond to replication stress by activating the replication stress checkpoint, which is initiated by ATR and Chk1 kinases (Branzei & Foiani, 2009). These kinases coordinate the protection of replication forks, the inhibition of late-origin firing, the prevention of cell cycle progression, as well as the induction of repair and the restarting of replication once the stress is resolved. Rif1 is a phosphatase involved in preventing late origin firing during replication stress, and consequently helps to maintain the established replication timing (Yamazaki *et al*, 2012; Foti *et al*, 2016). Once stress disappears, DNA synthesis may be reinitiated from the same arrested replication forks or, if the fork is collapsed, from the near-by backup licensed origins.

Oncogenes can induce replication stress and genomic instability, by reducing dNTP levels available for replication or by increasing conflicts between the replication and transcription machinery (Primo & Teixeira, 2020; Aird *et al*, 2013). Consistent with this, tumour cells exhibit higher levels of basal replication stress compared to non-transformed cells (Boyer *et al*, 2016; Ubhi & Brown, 2019; Nieto-Soler *et al*, 2016), and are highly dependent on replication stress response. Most probably, for this reason, mutations in ATR or Chk1 are uncommon in cancer cells, as many tumour cells have adapted to proliferate despite the presence of elevated replication stress. In fact, the response of tumour cells to a sustained replication stress differs from the one of non-transformed cells. Upon prolonged depletion of dNTPs induced by hydroxyurea (HU) treatment, non-transformed cells fail to reinitiate DNA replication after HU removal and instead enter senescence (Ercilla *et al*, 2016). This failure to restart DNA synthesis is due to two factors: the collapse of replication forks and the inability to efficiently activate new replication origins. The activation of APC/C^Cdh1^, which leads to Cyclin A degradation and thus results in the loss of CDK2/Cyclin A activity, has been previously identified as the cause of this incapability(Ercilla *et al*, 2016). In contrast, colorectal cancer (CRC) cells have been shown to reinitiate DNA synthesis and continue proliferating after recovery from prolonged replication stress. However, this process is accompanied by the acquisition and accumulation of DNA damage and is also associated with a lack of APC/C^Cdh1^ activation (Ercilla *et al*, 2016), although the underlying mechanisms remain unclear.

In this study, we investigated whether CRC cells rely on new origin firing to resume DNA synthesis after recovering from a severe replication stress, and if so, whether these origins are located in new replication domains. Given the critical role of origin activation in maintaining chromatin structure, we also explored how this process impacts chromatin accessibility, gene expression, and replication timing. Our findings revealed that DNA synthesis reinitiation following severe replication stress leads to lasting changes in CRC cells with an increase in chromatin accessibility, a significant dispersion of nuclear morphology, a shift in replication timing, and an alteration of gene expression such as *KRT20*, a developmentally regulated gene commonly used as marker of tumour differentiation. These alterations are pivotal in driving chromatin and genomic instability, providing new insights into the mechanisms that enable CRC cells to adapt and continue to proliferate under conditions of replication stress.

## RESULTS

### Origins fired upon recovery from a prolonged replication stress are spatially separated from stalled forks in CRC cells

In contrast to non-transformed hTERT-RPE cells, we previously demonstrated that HCT116 CRC cells can resume DNA synthesis when recovering from a severe and prolonged replication stress induced by a 14h treatment with 10 mM hydroxyurea (HU) Click or tap here to enter text.(Ercilla *et al*, 2016)Click or tap here to enter text.. Notably, HCT116 cells did not activate APC/C^Cdh1^, and consequently did not degrade Cyclin A upon this prolonged replication stress (Ercilla *et al*, 2016). These findings suggested that cancer cells may be able to trigger the activation of new origin during recovery to restore DNA synthesis. To test this hypothesis, we performed DNA fiber analysis. We first confirmed that after acute replication stress (2h HU) approximately 90% of replication forks were able to restart in HCT116 cells, similar to the recovery observed in control cells (Fig.1A-C). In contrast, following prolonged replication stress (14h HU) only 45% of replication forks restarted, and about 55% of the replication forks were stalled. Strikingly, a significant increase in new origin firing was observed under prolonged replication stress, with 40% of origins newly activated (Fig.1A-C). This increased activation of new origins after prolonged replication stress may facilitate cell cycle progression despite replication challenges (Sup Fig.1D). As expected, inhibition of CDK2 – a kinase known to promote origin firing - significantly reduced the activation of new origins from 32% to 4% (Fig.1D-F), slightly decreased HU-treated cell viability (Sup Fig.1A-B), and substantially increased the time required for the cells to complete DNA replication and progress to the G2 phase (Sup Fig.1C-D). To assess whether these newly activated origins were located within the same replication domain (RD) as the previously stalled replication forks, we performed an immunofluorescence analysis of replication foci. Replication foci were defined as 130 nm circles, as described in the Methods section and according to reference (Zhao *et al*, 2017). CellsClick or tap here to enter text.were sequentially labelled with two thymidine analogues: 5-chloro-2’deoxyuridine (CldU) for 15 min to mark the forks active before replication stress, and 5-iodo-2’deoxyuridine (IdU) to identify the forks that were reinitiating after HU removal (Fig.2A). From this pulse-chase experiment, we quantified the percentage of newly activated foci that colocalised with pre-existing ones (Fig.2A). Three controls were included to validate the foci colocalization settings: two for maximum colocalization and one for minimum colocalization (Fig.2A). Maximum colocalization was expected when cells were sequentially labelled with CldU and IdU without a release period ([C-I15’] and [C-I30’]). Of note, to ensure sufficient recovery time after prolonged replication stress, we used a 30-min IdU labelling period, as cells typically resumed DNA synthesis approximately 15 minutes post-stress (Feu *et al*, 2022a). Under non-stressed conditions, cells Click or tap here to enter text. with IdU immediately after CldU confirmed high colocalization with a median of 74% and 67% of foci co-labelled with both markers per cell for C-I15’ and C-I30’, respectively (Fig. 2A-C). Minimum colocalization was anticipated in cells with a 1h release in normal media between labels (Fig.2A, [C-1h-I]), as this interval would allow the activation of new RDs, thus resulting in non-colocalized foci (Fig.2A-B). Under this condition, we observed a median of 41% overlap between IdU- and CldU-labelled foci per cell (Fig.2B-C), representing a 30% decrease in colocalization compared to conditions of maximum colocalization. Notably, following severe replication stress, IdU-labelled foci showed a reduced colocalization with the CldU-positive foci with a median of 59 %. Interestingly, we showed that an average of 65% replication forks restarted under these conditions (Fig.1), which would typically appear as colocalization events. Thus, this decrease in foci colocalization indicates that most new origin firing during recovery from prolonged replication stress occurred in separate foci, spatially distinct from those of arrested forks (Fig.2B-C, [C-HU-I30’]). Our results suggest that the new origins firing after HU treatment are associated with different replication domains.

**Fig 1.**
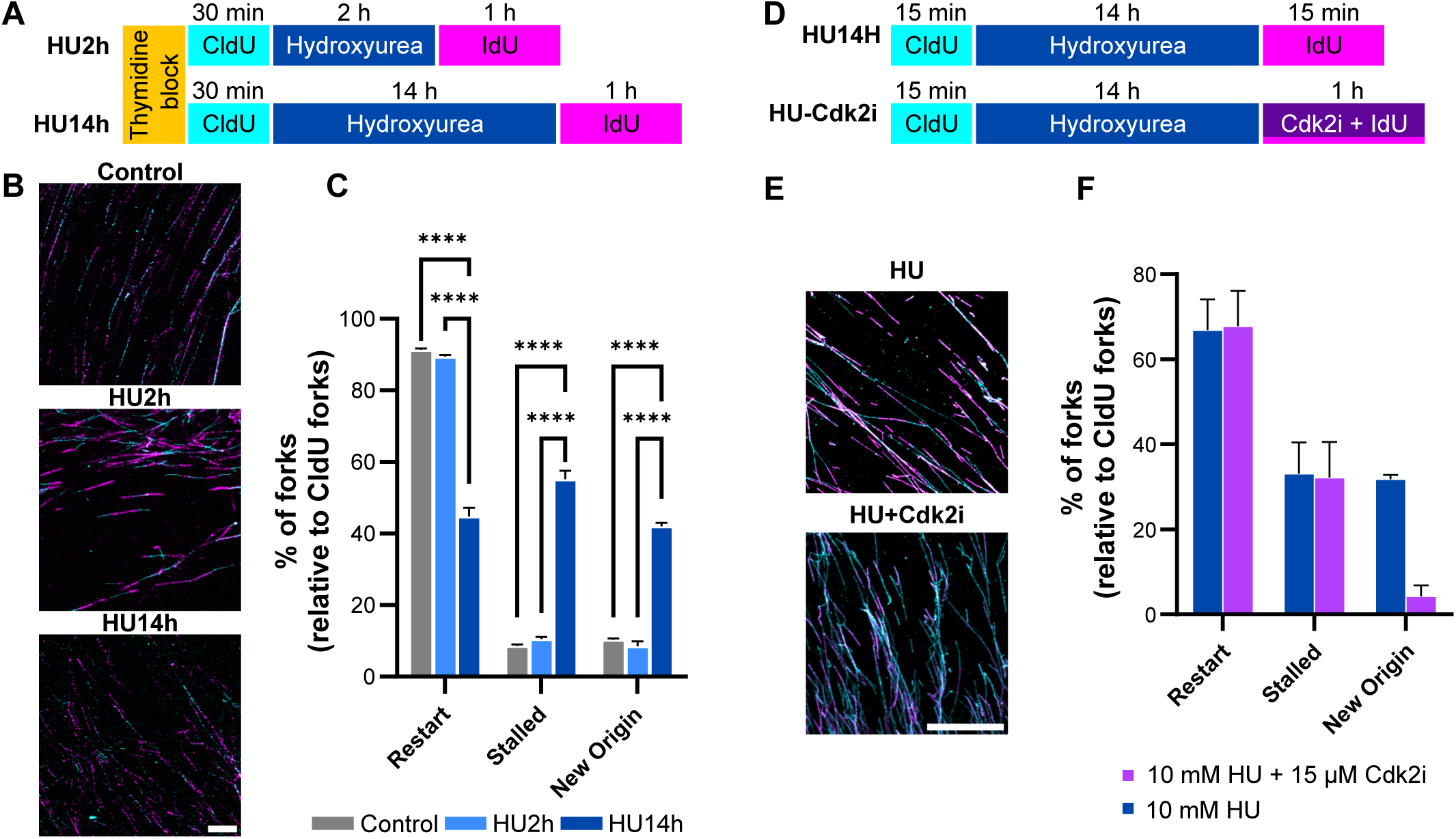
Recovery from HU-induced severe replication stress induced new origin activation. **A.** Schematic representation of the experimental design. HCT116 cells were synchronized in S phase using a single thymidine block. Cells were pulsed-chased with CldU, 5-chloro-2′-deoxyuridine, and IdU, 5-iodo-2’-deoxyuridine. Cells were treated with hydroxyurea (HU) for 2h or 14h to induce mild or severe replication stress, respectively. **B.** Representative images of DNA fiber assay under the experimental conditions of (A). DNA labelled with CldU are shown in cyan, IdU-stained DNA are shown in magenta. Scale bar, 10 µm. **C.** Quantification of replication fork dynamic (see methods). The percentage of replication fork restart, stalled replication forks and new origin firing events were quantified and normalized to the total of CldU -labelled fibers. Data were collected from over 1500 fibers per condition across three independent experiments. Results are presented as Mean ± S.E.M. Statistical analysis was conducted using paired t-test comparing each condition to control (C), with ****, p-value < 0.0001. **D.** Experimental design. Asynchronous HCT116 cells were labelled as indicated. Cells were treated with HU and CDK2i as outlined. HU, hydroxyurea; CldU, 5-chloro-2′-deoxyuridine; IdU, 5-iodo-2’-deoxyuridine; CDK2i, CDK2 inhibitor type II. **E.** Representative images of DNA fiber assay where fibers were stained with CldU (shown in cyan) and with IdU (shown in magenta) as indicated. Scale bar, 50 µm. **F.** Quantification of fork outcomes. Quantification of replication forks relative to the total of CldU-labelled fibers. Data were obtained from a minimum of 1000 fibers across two independent experiments. Results are presented as Mean ± S.E.M.

**Fig 2.**
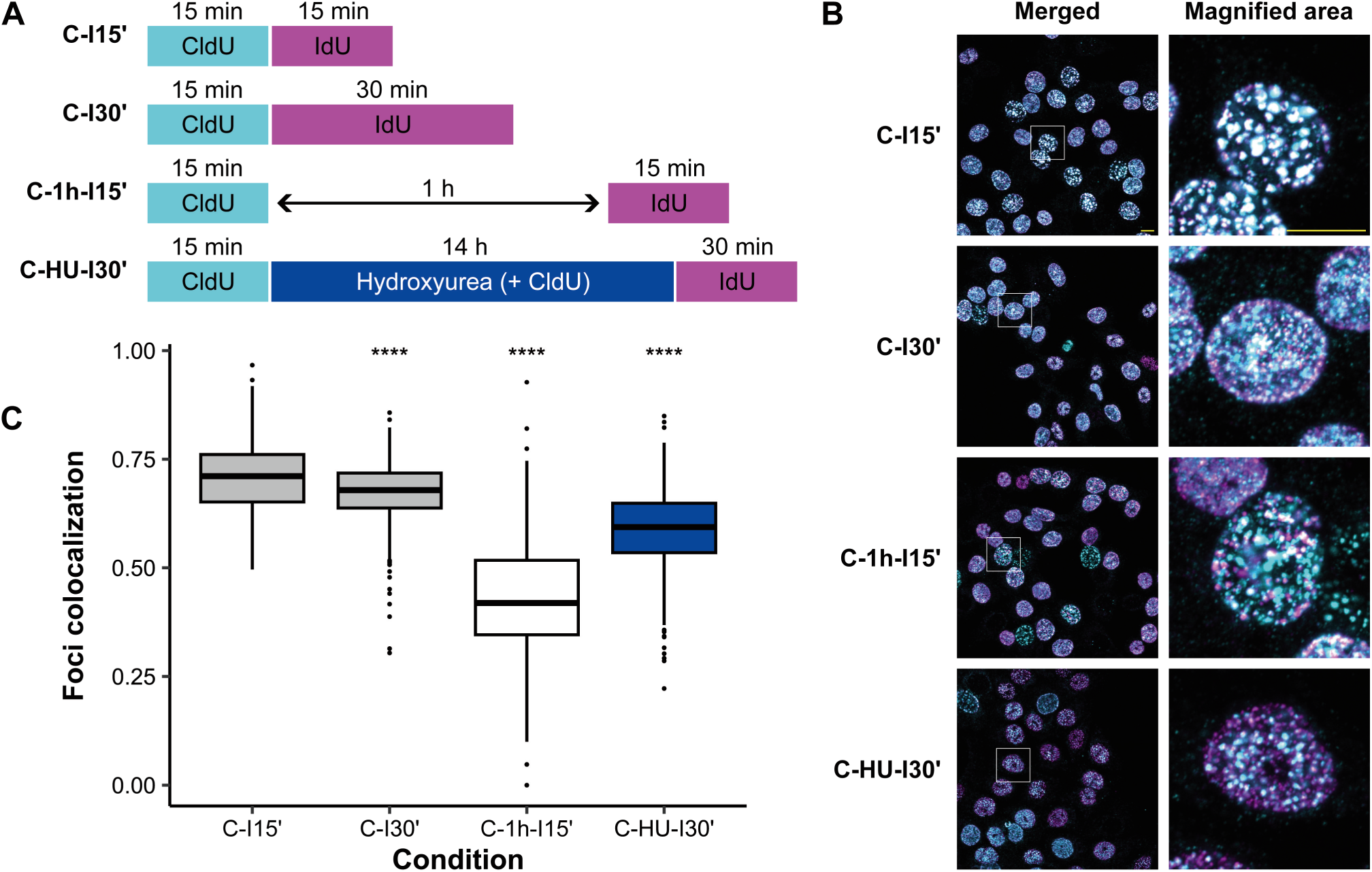
HU-induced replication stress disrupted replication dynamics in HCT116 cells. **A.** Experimental setup. Asynchronous HCT116 cells were treated and labelled as indicated. HU, hydroxyurea; CldU, 5-chloro-2′-deoxyuridine; IdU, 5-iodo-2’-deoxyuridine. **B.** Representative confocal images of replication foci. Immunofluorescence images show replication foci with CldU-labelled regions in cyan and IdU-labelled regions in magenta. Insets on the right show magnified nuclei. Scale bar, 100 µM. **C.** Quantitative analysis of replication foci colocalization. High-content screening quantification of replication foci colocalization was performed measuring the ratio of CldU-IdU positive foci to the total of IdU positive foci per nuclei. Image analysis was performed using ImageJ software, segmenting replication foci, and analysing the colocalization (see methods). Data are presented in a boxplot; the box edges represent the 25^th^ and the 75^th^ percentiles. The black line in the box denotes the median and whiskers show the range up to 1.5 IQR of the population. Significance is calculated by Wilcox test. Individual points indicate potential outliers. At least 1000 cells were analysed per condition (N=2).

### DNA synthesis reinitiation after severe replication stress induces durable changes in chromatin accessibly and nuclear size

We investigated whether the reinitiation of DNA replication in distinct RDs after replication stress could lead to alterations in chromatin compaction and structure, and whether these changes would persist into the next cell cycle. First, we determined that cells treated with HU for 14h entered mitosis upon release, with a pic between 10 and 13h post-treatment (Sup. Fig.2A-B). We further characterized that 80% of cell population had duplicated their DNA (EdU-positive cells) for a second time after 36h of release from HU treatment (Fig.3B-C), proving that cancer cells continued cycling even after having undergo a severe replication stress. Interestingly, during HU-treatment and recovery, we observed significant changes in nuclear size, with nuclei almost doubling in size after 48h of release (Fig.3D-E and Sup. Fig.3). Specifically, HU treatment resulted in a more heterogeneous nuclear size distribution among cells, and remarkably, these alterations persisted for at least 48h post-release. By this time, cells had already entered the next cell cycle phases and the median nuclear size was 91 (IQR: 76 – 106) in control conditions and 133 (IQR: 107 - 162) after 48h of recovery (Fig.3D-E). These findings prompted us to analyse if chromatin accessibility was modified. To this end, Assay for Transposase-Accessible Chromatin using sequencing (ATAC-seq) analysis was performed on control cycling cells (C), cells treated for 14h with 10 mM HU, and cells that recovered for either 12h (HUR12) or 36h (HUR36) post-HU treatment (Fig.4A). A total of 165315 accessible regions were identified across all conditions, with varying numbers of peaks detected in each (between 64957 and 88335 for each). The number of accessible regions in each one of the intersections between conditions is shown in figure 4B. We specifically focused the analysis on regions that became newly accessible or that lost accessibility during recovery, by evaluating only the common regions between the experimental conditions HUR12 and HUR36 that were absent in control (C) and HU-treated cells (Fig.4B, column V), and *vice versa* (Fig.4B, column III). We found that after recovering from severe replication stress, 2576 regions gained accessibility (Fig.4B, column V), while accessibility was lost in 3975 regions (Fig.4B, column III).

**Fig 3.**
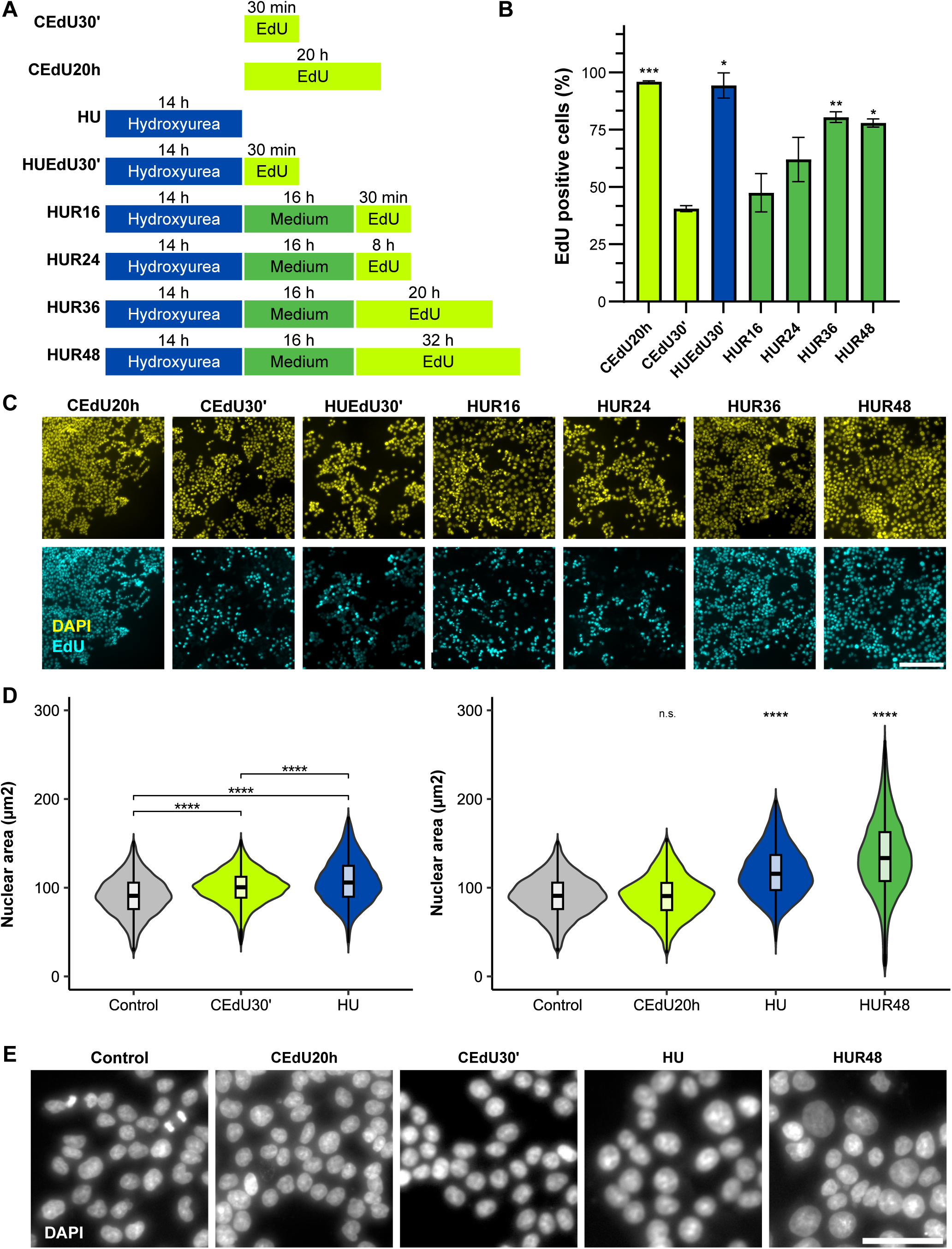
HCT116 cells resume cell cycle and arrive to a second S-phase after HU-induced replication stress with an increase in the nuclear area. **A.** Experimental design. Schematic representation of the experimental setup, using, 5-ehynyl-2’deoxyuridine (EdU) to label DNA synthesis and hydroxyurea to induce replication stress. **B.** Quantification of EdU incorporation after first mitosis during HU release. Proportion of cells incorporating EdU after progressing through mitosis upon release from HU-induced replication stress (also see Sup. Fig.3). Data represent mean ± S.E.M from three independent experiments (N=3; also see Sup. Fig.4). Statistical analysis was performed using ANOVA with multiple comparisons in GraphPad. Significance levels are shown relative to CEdU30’. *, p < 0.05; **, p < 0.01; ***, p < 0.001. **C.** Representative images of EdU incorporation (in cyan) in each experimental condition. Nuclei were counterstained with DAPI (in yellow). Scale bar, 100 µm. **D.** Analysis of the nuclear area. Nuclear area analysis in cells recovering from HU-induced replication stress was conducted from images with more than 5,000 cells of three independent experiments. Data are presented as a boxplot. The box represent mean, 25^th^ and the 75^th^ percentiles with an overlaying violin plot to display nuclear area distribution. Statistical analysis was performed using a Kruskal-Wallis’ test (non-parametric ANOVA) followed by Dunn’s test for multiple comparison. Significance levels with comparisons made relative to control condition are denoted by: *, p < 0.05; **, p < 0.01; ***, p < 0.001; ****, p < 0.0001 and n.s., not significant. **E.** Representative images of DAPI-stained nuclei for each indicated experimental conditions. DNA is shown in grayscale. Scale bar, 50 µm.

**Fig 4.**
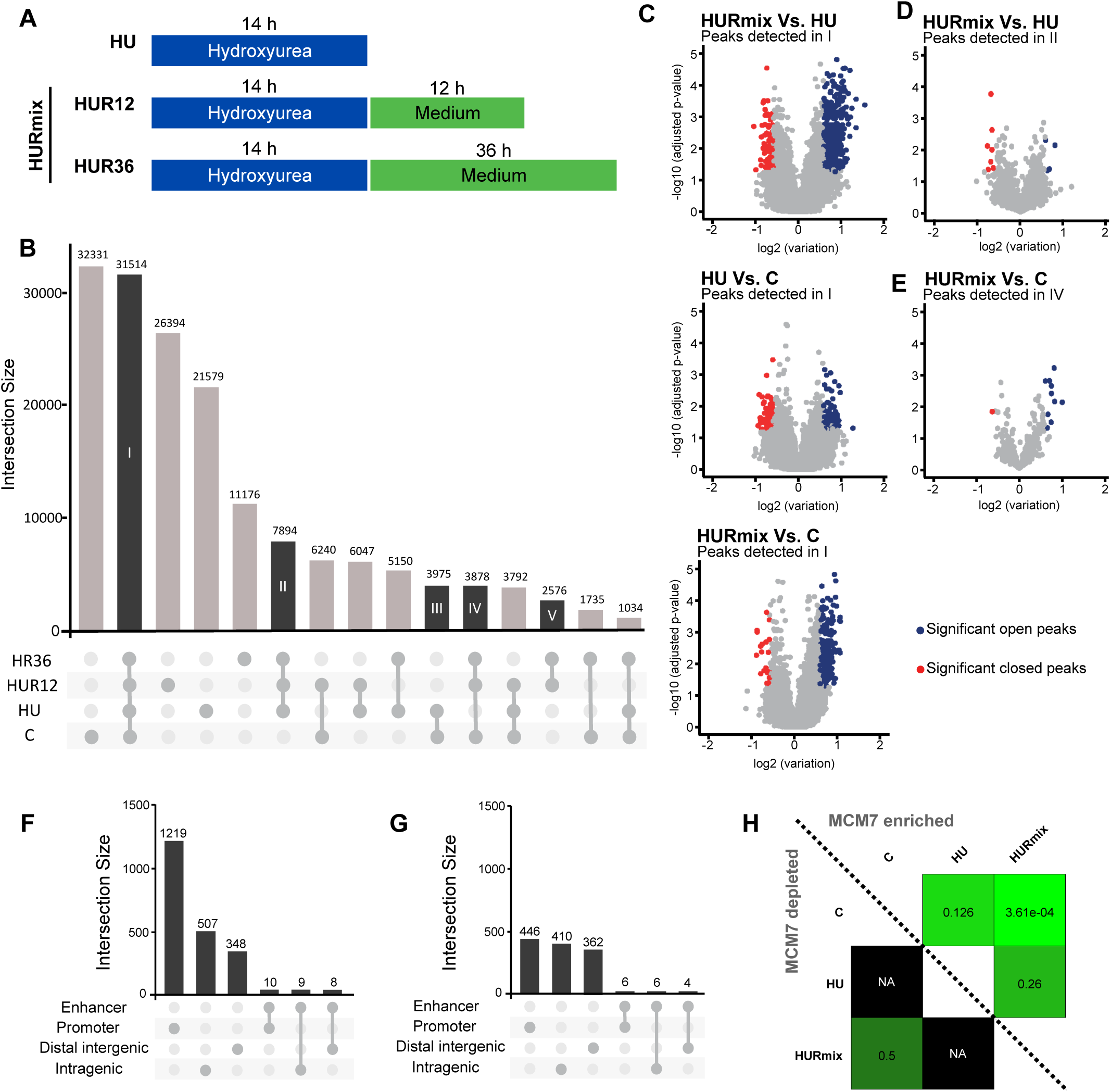
Chromatin accessibility is increased during HU-induced replication stress recovery in HCT116 cell line. **A.** Experimental design. Schematic overview of the experimental setup. Cycling cells correspond to control, presented as C. Cells were treated with hydroxyurea (HU) and subsequently released (R) before processing for ATAC-seq analysis. Release conditions analysed as a single combined condition are named “HURmix”. **B.** ATAC-seq data analysis. Graphical representation of the peaks detected across three independent experiments. Column bars represent the size of peak intersections (number of peaks) for each condition or condition group (indicated below), with the number of peaks detected indicated above each column. Column I shows peaks common to all samples. Column II shows peaks present during HU treatment and maintained during the recovery period. Column III represents peaks lost during replication stress recovery. Column IV displays peaks lost only during HU treatment, and column V contains peaks detected exclusively in the release condition. **C-E**. Volcano plots of ATAC-seq peaks across conditions. Volcano plots display peaks detected in at least in two of three experiments of ATAC-seq. Peaks with significative increase are shown in blue, and those with significant decreases in red. Panel (C) highlights the analysis of the peaks detected in all the conditions (Column I in panel B). Panel (D) represent the analysis of peaks present during HU treatment and recovery period (column II in panel B). Panel (E) shows peaks detected in control and recovery conditions (column IV in panel B). **F.** Genomic localization of gained peaks. Representation of the distribution of peaks identified as gains (column V of panel B) and significant increases (blue dots in C, D and E) across genomic regions including enhancers, promoters, distal intergenic and intragenic regions. Column bars show the peak intersection sizes (number of peaks). **G.** Genomic localization of lost peaks. As in panel (F), the distribution of lost peaks (column III in panel B) and significantly decreased peaks (red dots in panels C, D and E) across genomic regions. **H.** Enrichment analysis of MCM7-associated peaks. Heatmap representing enrichment analysis of ATAC-seq peaks overlapping MCM7 regions, with enrichment levels indicated on a green gradient scale for each comparison. The top-right diagonal shows increase in enrichment, while bottom-left diagonal shows decrease. Corresponding p-values are showed in each condition.

In addition, we analysed quantitative changes in the accessibility of regions detected across all conditions and assessed the percentage of cells presenting an increase or a decrease of accessibility within these accessible regions (Fig.4B, column I). To facilitate the analysis, we combined the two recovery conditions (HUR12 and HUR36) into a single category called HURmix (Fig.4A). The fold changes and the statistical significance of these alterations in chromatin accessibility are depicted in the volcano plots (Fig.4C-E). Notably, when comparing the recovery conditions (HURMix) to control cells or to cells treated with HU without recovery (HU), there were more gains in accessibility than losses (Fig.4C). In contrast, the comparison between HU-treated and control cells showed less pronounced differences, with no clear predominance of increases in accessibility (Fig.4C). Analysis of gained or lost peaks showed that only few of them exhibited significant changes (Fig.4D-E). Indeed, the most significant changes were observed in peaks consistently detected in all the conditions (Fig.4B, column I), rather than in peaks that are entirely gained or lost (Fig.4B, column III and V).

We further classified the detected regions based on their location and their surrounding gene regulatory context, categorizing them into four groups: promoter, enhancer, intragenic and distal intergenic regions. Additionally, we analysed changes in known enhancer regions, which could belong to any of these four categories. After recovering from severe replication stress, we observed a specific increase in accessibility in promoter regions (Fig.4F). In contrast, the regions exhibiting loss or decreased accessibility showed no preference among the categories (Fig.4G). Considering that the total size of the promoter regions is significantly smaller than that of intergenic or intragenic regions, our data suggest that the reinitiation of DNA synthesis upon replication stress induces specific changes in promoter accessibility. Interestingly, in some cases this increase in accessibility was maintained during the next cell cycle. The increased accessibility in promoter regions aligns with the current understanding that active promoters are preferential sites for DNA replication initiation (Akerman *et al*, 2020; Cayrou *et al*, 2015; Jodkowska *et al*, 2022; Sequeira-Mendes *et al*, 2009). Examples of genes with increased chromatin accessibility in their promoter or enhancer regions are provided (Sup.Fig.4). Interestingly, the enhancer region of the *KRT20* gene also exhibited increased accessibility during recovery, although this peak was not initially detected with our peak calling algorithm, possibly due to the proximity of a larger *KRT23* enhancer peak (Sup.Fig.4A). This finding indicates that additional regions than the ones we have detected could be changing the accessibility. Gene ontology analysis performed on all the promoter regions with altered accessibility, comparing control and HUR48 conditions, did not reveal a pronounced enrichment of any specific gene set, thus suggesting that the changes detected in chromatin accessibility were not specifically directed towards activating a particular subset of genes.

If the new accessible regions resulted from the activation of new origins during the recovery from severe replication stress, we would expect them to be enriched in MCM binding regions. A large-scale analysis of MCM7 DNA binding regions reported in ENCODE 3 (Transcription Factor ChIP-seq E3, UCSC genome browser, hg19) (Dunham *et al*, 2012) served as a base for our analysis of possible MCM7 peaks enrichment in the regions differentially accessible in our ATAC-seq dataset. The analysis revealed that MCM7 DNA binding sites shared 1113 peaks with our overall ATAC-seq peaks, with most of them (656) located in regions conserved across conditions (p-value=e-182, FC=3.0). We considered the peaks that appeared on the whole set (31514 peaks), and merged the MCM7 peaks with differential peaks from the comparisons of HURmix vs C, HURmix vs HU, and HURmix plus HU vs C. While most peaks remained conserved across conditions, a significant number of peaks that gain accessibility post-release (HURmix) were located on MCM7 binding sites, reinforcing the connection between chromatin accessibility changes and DNA replication reinitiation following replication stress (Fig.4H).

### Persistent changes in replication timing and gene expression are triggered by DNA synthesis reinitiation following severe replication stress

We investigated whether the activation of DNA replication origins in new foci, distinct from those active before replication stress (Fig.1 and Fig.2), could lead to persistent changes in DNA replication timing. To this end, the replication timing of genes of interest was analysed in control cells and compared to the replication timing of cells exposed to severe replication stress (10 mM HU for 14h) and then allowed to recover for 48h (Fig.5A). This recovery period was chosen to ensure that cells had completed an additional DNA replication cycle after the release from replication stress (Sup.Fig.2). Since we hypothesized that the newly activated origins might overlap with regions showing increased chromatin accessibility during replication stress recovery, we focused our analysis on genes in these regions, such as *BMP1*, *FA2H*, *GAPDH*, *HBA*, *HBB*, *KRT20*, *KRT23* and *NETO1* (Fig.5B-C; Supplementary figure 4). We captured replicating DNA by BrdU incorporation and immunoprecipitation. Our results showed that, for instance, the regions of *HBA* and *NETO1* genes that are replicated in the late S-phase in control conditions, are activated prematurely in mid-S-phase after HU-induced replication stress (Fig.5C). Conversely, replication timing was delayed upon HU treatment in some cases such as *BMP1*. Normally active in early S-phase under control conditions, this region was activated and detected only later in mid-S phase post-HU treatment (Fig.5C), thus suggesting that this region collapsed after HU-induced replication stress and required additional time to be replicated. We propose that this delay might be due to the firing of flexible origins flacking the collapsed region to aid recovery, which is in agreement with our previous demonstration that the new fired origins were located in different foci (Fig.2). Importantly, our results demonstrated that a significant replication timing shift also occurred in cells 48h post-HU recovery compared with untreated cycling cells, indicating that severe and prolonged replication stress produced durable changes in replication timing (Fig.5B-C). When comparing the replication timing 48h after release from HU treatment to control conditions, we observed some regions where replication timing did not fully revert to their original replication timing patterns. For instance, *NETO1 and HBA* showed advanced replication timing during recovery, corresponding to an early replication activity (Fig.5C). Conversely, *KRT23* initially replicated earlier after HU treatment, but shifted to later S-phase replication timing during recovery (Fig.5C). Some other genes, such as *BMP1*, that already exhibited delays in replication activity after HU treatment, presented a more exaggerated delay during the recovery (Fig.5C). Overall, most of the regions studied presented a shift in replication timing program following HU-induced replication stress, with this shift becoming even more pronounced during 48h post-(Hiratani *et al*, 2008)recovery period, and with early-to-late timing shifts being more prevalent than late-to-early shifts.

**Fig 5.**
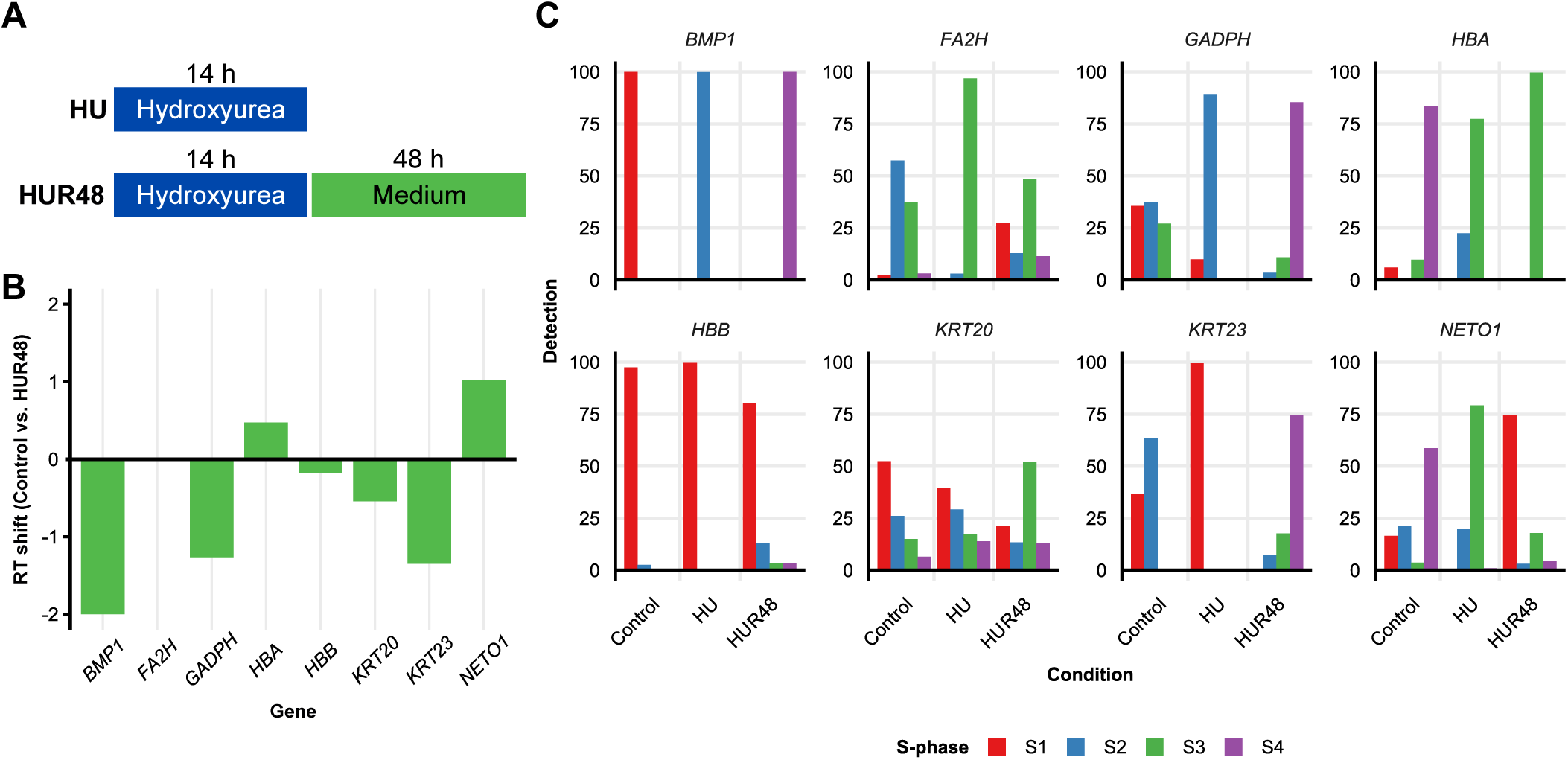
Replication timing (RT) is modified after HU-induced replication stress. **A.** Experimental design. HCT116 cells were treated with hydroxyurea (HU) to induce replication stress and further released (R) for recovery for 48h R. **B.** Replication timing (RT) shift correspond to the comparison between cells recovered for 48h post-HU treatment (HUR48) and untreated cycling cells. RT shift was calculated as indicated in methods. Positive values indicate an advance in RT (shift from late to early S phase), while negative values correspond to a delay in RT (shift from early to late S phase). **C.** Analysis of nascent DNA strands. qPCR results of selected genes across four fractions of the S phase. The results were normalized to the total detection in all fractions (S1+S2+S3+S4). Fractions are defined as followed: S1, early-early S phase (in red); S2, early-mid S phase (in blue); S3, mid-late S phase (in green); S4, late-late S phase (in purple).

As changes in chromatin accessibility and replication timing have been directly related to changes in gene expression (Sequeira-Mendes *et al*, 2009), we next assessed whether replication stress could permanently alter gene expression. To this end, we studied the expression of *KRT20*, (Fig.6A and Sup.Fig.5), *KRT23*, *SPARC*, *TOP2A* and *FA2H* (Sup.Fig.5), as these genes have enhancers, promoter or transcription start-sites displaying a significant increase in chromatin accessibility upon recovery from HU treatment. Of note, *KRT20* and *KRT23* are developmentally regulated genes commonly used as markers of tumour differentiation (Nakatani *et al*, 2024; Hiratani *et al*, 2008). Following recovery from HU treatment, *KRT20, SPARC* and *KRT23* expression levels increased and persisted up to 48h post-HU removal (Fig.6B-D and Sup.Fig.5). Notably, this increase in expression was reduced in the presence of CDK2 type II inhibitor, which prevents DNA replication initiation from new origins (Sup. Fig.5). In contrast, other genes with increased promoter accessibility, showed no change in expression, thus suggesting that additional factors beyond chromatin accessibility may be necessary for transcription activation.

**Fig 6.**
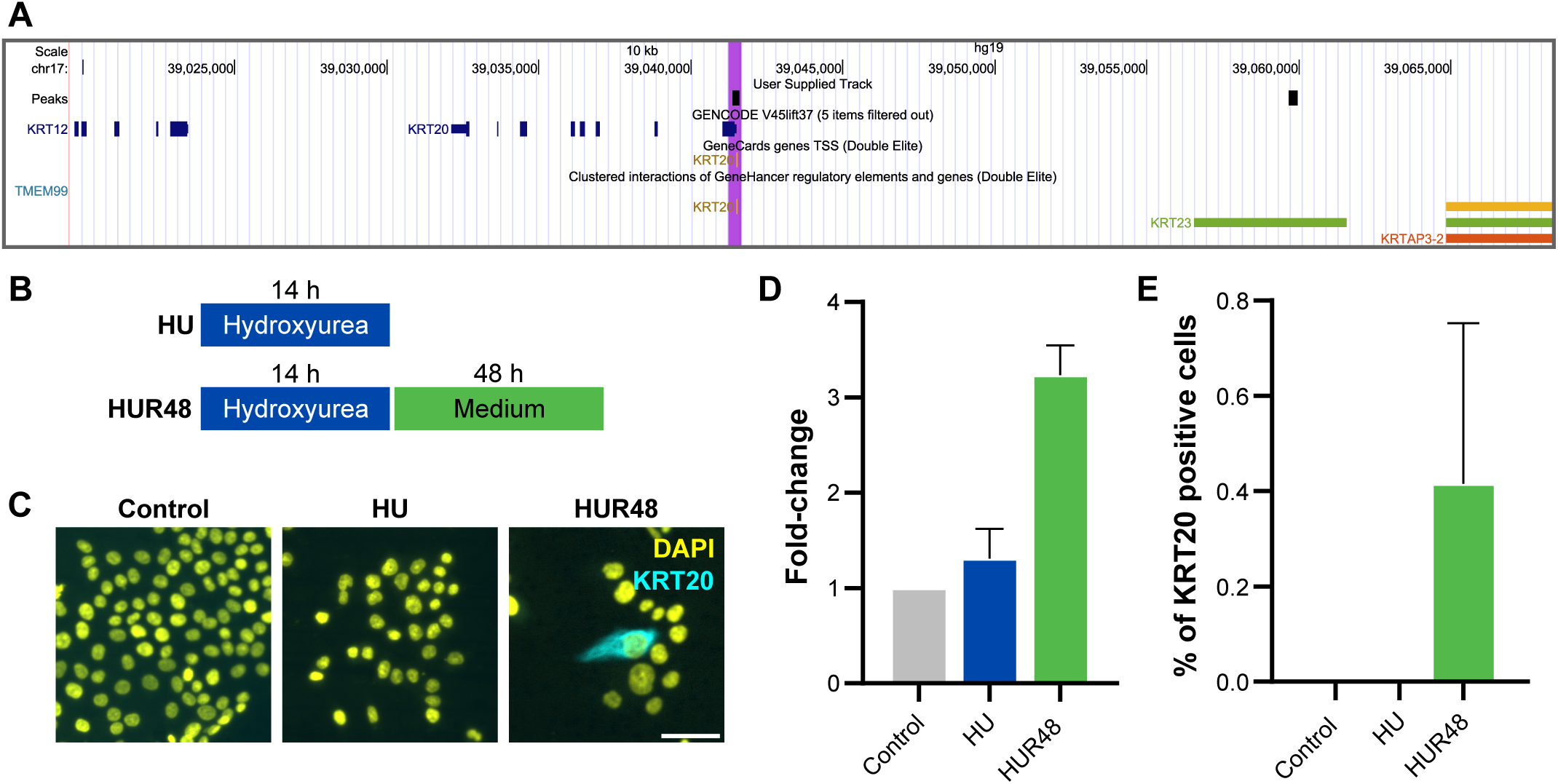
Transcription is altered during the recovery of HU-induced replication stress in HCT116 cell line. **A.** UCSC Genome Browser visualisation of the KRT20 genomic region with four tracks. The first track highlights the peaks detected in ATAC-seq analysis (see Fig. 4). The second track shows GENCODE genes annotations. The third track presents transcriptional start sites (TSS) from GeneCards. The fourth track shows the clusters of genetic interactions from GenHancer. **B.** Schematic representation of the experimental design. HCT116 cells were treated with hydroxyurea (HU) to induce replication stress. **C.** Representative images of CK20-positive cells. Immunofluorescence was conducted to detect endogenous CK20 (encoded by *KRT20*) in cyan. Nuclei were counterstained with DAPI (in yellow). Scale bar, 50 µm. **D.** qPCR analysis of *KRT20* expression. Data is presented as the fold change in *KRT20* mRNA levels compared control cells and cells released 48h-post HU treatment. Data represent the mean of four independent experiments ± S.E.M. **E.** Quantification of CK20-positive cells in optical images. Data show the number of CK20-positive cells normalized total number of cells in 2 independent experiments. Data is presented as the mean of both experiments ± S.E.M.

Finally, *KRT20* protein expression (CK20) was analysed by immunofluorescence. While almost absent in control and HU treated cells, CK20-positive cells were detected at 48h HU post-release (Fig.6C-E). Immunofluorescence data revealed that the increased in CK20 expression was not faint and homogenous among all cells, but instead occurred only in a subset of cells with strong staining, indicating a stochastic activation of *KRT20* likely triggered during HU release rather than a programmed response to replication stress (Fig.6C-E).

Overall, our findings indicate that CRC cells can bypass replication stress responses by activating new replication origins, altering replication timing, and modifying gene expression patterns. This adaptive capability allows CRC cells to continue proliferating even as they accumulate epigenetic instability, a factor known to accelerate tumour progression and heterogeneity. This study sheds light on the enduring consequences of the ability of cancer cells to tolerate high levels of replication stress and provides a better understanding of cancer cell biology to potentially open new avenues for developing therapies aimed at targeting replication origin activity to weaken cancer cell resilience.

**Fig 7.**
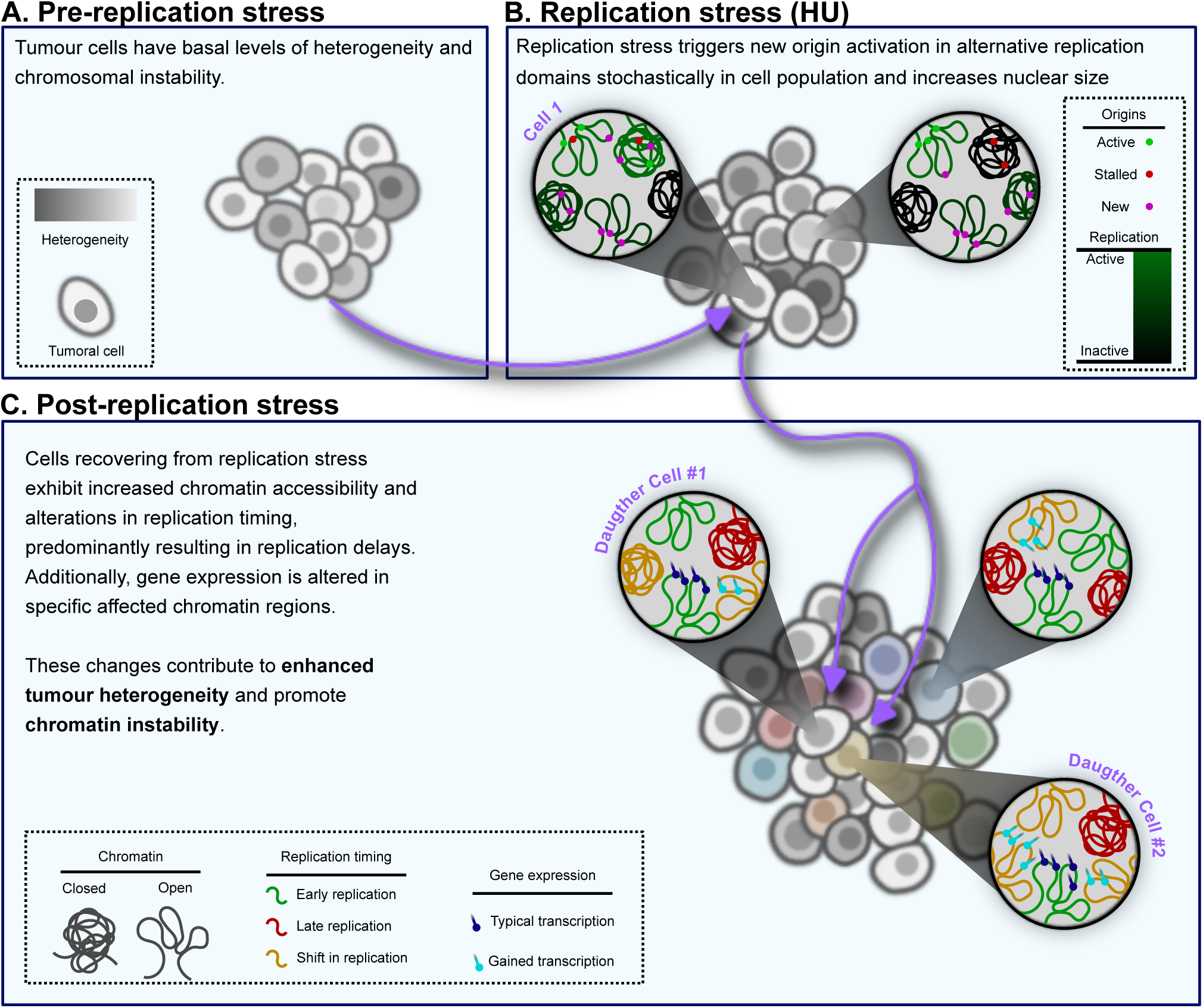
Schematic representation of our proposed model for colorectal cancer cell adaptation to replication stress. This model illustrates the dynamic response of colorectal cancer cells to replication stress. During pre-replication stress, cells exhibit basal levels of heterogeneity and chromatin stability. However, during replication stress, cells encounter replication challenges that trigger stochastic activation of new replication origins within alternative replication domains. At this stage, cells also showed an increase in nuclear size, which reflects structural and functional adaptations to stress. In the post-replication stress recovery phase, cells that successfully recover display an increased chromatin accessibility, particularly in promoter regions, alongside with alteration of replication timing, predominately delays. These changes are associated with gene expression alterations, including sustained transcriptional shift in some genes, which contribute to their ability to adapt to stress. Together, these adaptive mechanisms promote tumour heterogeneity and chromatin instability, thus highlighting the role of replication stress recovery in driving cancer cell resilience and evolution.

## DISCUSSION

It is now accepted that not only genomic instability, but also epigenetic instability are hallmarks of cancer cells (Hanahan, 2022). Both types of instability are key drivers of increased tumour heterogenicity aiding the adaptation of tumour cells to environmental pressures, thereby promoting tumour development and resistance to treatments. While the mechanisms underlying genomic instability are well-studied (Burrell *et al*, 2013b), the cellular processes leading to chromatin instability remain unclear (Hsu *et al*, 2021; Jasencakova *et al*, 2010).

Replication stress is intrinsic of most tumour cells and is known to promote genomic instability even in the absence of external mutagenic agents (Burrell *et al*, 2013b; Gaillard *et al*, 2015). Although histone modifications under replication stress are well-documented (Jasencakova *et al*, 2010), few studies explored permanent changes in chromatin activity (Courtot *et al*, 2021). Here, our findings strongly support the hypothesis that replication stress induces stochastic changes in chromatin accessibility and activity in cancer cells, and most importantly, that these changes persist through subsequent cell cycles, thus contributing to long-term epigenetic variability and increased cell population heterogenicity. Our results also indicate that activation of new replication origins in distinct RD are key elements that drive these changes.

Interestingly, this mechanism may be a specific feature of tumour cells. Under mild replication stress, DNA replication can recover in both non-transformed cell lines (Ercilla *et al*, 2016) and in tumour cells (Fig.1) through replication fork restart and activation of adjacent dormant origins. However, non-transformed cells exposed to severe and prolonged replication stress typically do not recover DNA replication, and instead activate APC/C^Cdh1^ and enter senescence (Ercilla *et al*, 2016). Conversely, under similar severe and prolonged stress, CRC cells can reinitiate DNA synthesis. Here, we showed that CRC cells recover DNA replication by activating new origins. Most importantly, these new origins are located in different replication foci, rather than dormant origins.

In somatic cells, not all origins are activated simultaneously, instead they are triggered in groups according to a precise and robust replication timing, which is characteristic for each cell lineages to maintain cell identity (Pope *et al*, 2014; Rivera-Mulia & Gilbert, 2016b; Ryba *et al*, 2010; Wilson *et al*, 2016). However, replication timing initiation program exhibits remarkable flexibility in metazoan cells, and not all the cell population activates exactly the same origins, and the frequencies of specific origin initiation is distinct in different cell lineages (Smith *et al*, 2016; Cayrou *et al*, 2015). In our study, HCT116 cells recovering from prolonged HU treatment activated origins located in different loci that might be either late constitutive origins – the ones that would have been fired at the end of a normal S-phase -, or flexible origins – origins that would not have been used under normal conditions.

The observed initiation of DNA replication in new foci indicates the activation of new replication domains (Rivera-Mulia & Gilbert, 2016b; Wilson *et al*, 2016). Furthermore, we demonstrated that these changes are not transient but persist through the next cell cycle. While we anticipated primarily earlier replication timing of late origins, the analysis of S-phase fractions for 8 genes revealed varied shifts: two genes advanced in replication timing, four genes delayed replication, and two showed increased replication timing heterogenicity in the cell population.

Remarkably, ATAC-seq analysis revealed that recovery from severe and prolonged replication stress induced profound and sustained changes in chromatin accessibility, especially in promoter regions. During recovery from replication stress, we observed mainly gains in chromatin accessibility, and most of the changes were increases (and not complete gains). Thus, our results indicate that these sites were initially accessible in a low percentage of the cell population and that after replication stress, these specific sites became accessible in a higher percentage of cells. The fact that these changes were preferentially observed in promoter regions could suggest the activation of genes involved in stress response, however GO analysis did not confirm specific gene activation. Interestingly, replication origins have been shown to be associated with promoter regions (Jodkowska *et al*, 2022; Sequeira-Mendes *et al*, 2009), thus our results would propose that this increase in accessibility could be related to an increase of the number of potential origins located in these regions. Again, it is important to remark that most of the changes were maintained until the next cell cycle - when cells had replicated again the DNA - hence that they were not due to the activation of transient replication stress responses. Using well-known MCM7 binding sites as a surrogated marker of DNA replication origin sites (Sugimoto *et al*, 2018), we found that regions with increased accessibility post-replication stress had a high incidence of MCM7 binding sites, thus supporting the hypothesis that a new origin activation is tied to regions with increased chromatin accessibility.

Chromatin condensation and structure significantly influence replication timing, with early RD linked to less condensed active chromatin, while late RD are associated with more condensed and repressive chromatin (Aladjem & Redon, 2017; Hiratani *et al*, 2008). However, while epigenetic marks have been shown to influence replication timing, replication timing can reversely influence epigenetic marks such as levels of chromatin acetylation (Lande-Diner *et al*, 2009). In agreement with that, we demonstrated here the presence of durable changes in gene expression alongside changes in RD activation, chromatin accessibility and replication timing. To analyse changes in gene expression, we selected genes which showed changes of accessibility in their promoter or enhancer regions after replication stress recovery as well as genes known to be developmentally regulated in the lineage of colon epithelial cells, since lineage-specific genes are known to belong to chromatin compartments where their replication timing is modified during development (Nakatani *et al*, 2024). While the correlation between increased chromatin accessibility and gene expression was modest and only clearly observed in one of the five genes analysed, our findings with *KRT20* support that prolonged replication stress can induce permanent changes in gene expression. The lack of expression changes in the other genes analysed could be due to the fact that, in addition to changes in chromatin accessibility, other factors are required to fully activate the transcription program. Interestingly, the expression of CK20 was only observed in a subset of cells, proving a stochastic event most probably driven by the activation of flexible origins in the *KRT20*-containing rather than a programmed activation directed by the stress response. Although single-cell analysis would be needed to further support this hypothesis, the fact that inhibition of new origin firing by CDK2 inhibition prevented *KRT20* expression strongly reinforces this working model.

In summary, our findings reveal that CRC cells adapt to prolonged replication stress and recover by stochastically activating origins located in new RDs. This results in permanent changes in replication timing, chromatin accessibility and gene expression in a subset of the cell population, thus participating to an increased epigenetic heterogenicity. Although extreme and prolonged replication stress may not commonly occur during tumour development, localized chromatin replication challenges could trigger similar responses under mild stress, for instance in specific regions where the chromatin is difficult to replicate or where replication forks would collapse even under mild stress conditions. In fact, a previous study using mild replication stress suggested that the few permanent replication timing changes occurred predominantly in tumour cells, often closely associated with common fragile sites (CFS) (Courtot *et al*, 2021). Our findings have potential therapeutic implications, as most CRC therapies commonly induce strong replication stress. Our results alert that new epigenetic marks and features, potentially favouring treatment resistance, may emerge in surviving cells. Thus, it would be important for future research to explore and design combination therapies to target and block new RD activation or senescence induction under replication stress to mitigate the development of resistance to anti-cancer treatment.

## MATERIAL AND METHODS

### Cell culture and drugs

The colorectal carcinoma cell line (HCT116) was purchased from Horizon Discovery Ldt. (Cambridge, United Kingdom), and was grown in Dulbecco’s modified Eagle’s medium (DMEM): HAM’s F12 (1:1) with stable glutamine and pyruvate acid (Biowest, BWSTL0103-500 and L0135-500, respectively) supplemented with 7% FBS (D016, Diagnovum). Hydroxyurea (H8627, Sigma) 10 mM was used to induce replication stress through the inhibition of ribonucleoside reductase. CDK2 type II inhibitor (221409, Santa Cruz) 15 µM was used to inhibit new origin firing. 5-Bromo-2′-deoxyuridine (BrdU, 10 µM; B5002, Sigma), 5-Chloro-2′-deoxyuridine (CldU, 25 µM; C6891, Sigma), and 5-Iodo-2′-deoxyuridine (IdU, 250 µM; I7125, Sigma) are thymidine analogues used to label replicating DNA. 5-etinil-2’-desoxiuridina (EdU; A10044, ThermoFisher) is a small thymidine analogue that can be detected using click chemistry, eliminating the need for DNA denaturation. Thymidine (2.5 mM; T1895, Sigma-Aldrich) has been used to arrest cells in S phase.

### DNA Fiber assay

DNA fiber assay was performed following the protocol described in [42]. Anti-BrdU antibodies were used to label CldU (Abcam; ab6326; 1/1000) and IdU (Becton Dickinson, Franklin Lakes, NJ, USA; 347580; 1/200). Images were acquired with confocal microscopes Leica TCS-SL or Zeiss LSM880 with a PLAN APO 63× oil immersion objective (numerical aperture 1.4). Images were analysed using Fiji software. The number of fibers analysed in each experiment is indicated in the corresponding figure legend.

### Immunofluorescence

For foci analysis cells were grown onto glass coverslips and labelled with thymidine analogues (CldU and IdU). Cells were fixed with 70% ethanol for 10 minutes at room temperature. Then, cells were denaturalized and permeabilized with denaturalization buffer (2 M HCl; 0.2% Triton X-100 in PBS) for 30 minutes at room temperature. Denaturalization buffer was washed 3 times with buffer borat (100 mM Na2BO4O7 equilibrated to pH 8.5 with 100 mM H3BO3). Cells were washed twice with PBS-T (1% Tween) and then incubated 30 minutes with 1% BSA. Primary antibodies against CldU and IdU were incubated for 1h at 37°C and secondary antibodies were incubated 45 minutes at 37°C. Coverslips were incubated with DAPI (D9564, Sigma-Aldrich) and mounted with Mowiol (81381, Sigma-Aldrich). Images were obtained with confocal microscope ZEISS LSM880.

To analyse cell and nuclear area, cells were labelled with EdU. Coverslips were washed thrice with PBS and fixed with PFA 4% for 10 minutes (PFA 37% diluted in PBS; 1.03999.1000, Sigma-Aldrich). After 3 washed with PBS cells were permeabilized with permeabilization buffer (20 mM glycine, 1% Triton X-100 in PBS). Click reaction was used to detect EdU labelling by incubating the cells with a click buffer (100 mM Tris-HCl pH8; 2 mM CuSO4; 1 M Alexa488-Azide – Invitrogen AI0266; 100 mM Ascorbic acid; added to ddH2O in this order) for 30 minutes on a rocker in the dark. After 3 PBS washes, cells were incubated in blocking buffer (20 mM glycine; 0.01% Triton X-100; 1% BS; in PBS) for 30 minutes in movement. DAPI staining was performed for 15 minutes and coverslips were mounted with Mowiol (81381, Sigma-Aldrich). Images were obtained in Widefield Leica AF6000 optical microscope.

To analyse CK20 expression coverslips were washed thrice with PBS and fixed with PFA 4%. After 3 washed with PBS cells were permeabilized 10 minutes and blocked for 30 minutes. Primary antibody against CK20 (13063S, Cell Signaling Technologies) and secondary antibody were incubated for 1h at room temperature. HCS Cell Mask (H32721) was used at 1:2500 dilution and stained for 30 minutes. DAPI and Mowiol mounting was done as explained above.

### Image analysis

Foci analysis was performed automatically using an in-house program in ImageJ as follows. For each IdU-positive focus in each cell, a 130 nm region was marked, representing the estimated replication domain size. For the first thymidine analogue introduced (CldU) the marked region was doubled in size to be more restrictive and to increase specificity in identifying non-colocalization. This adjustment ensured that if the IdU signal was adjacent to or overlapping with CldU, it was considered part of the same replication domain. The data obtained were processed in R studio.

For area and EdU incorporation analysis, we used automated analysis with ImageJ plugin. Nuclear areas were first segmented using StarDist plugin, then nuclear area and DAPI intensity were measured. Nuclear ROIs were saved for future analysis. Next, an automatic threshold for EdU intensity was applied, and EdU signal was measured within the nuclear ROIs. The results were subsequently analysed with R studio.

### Replication timing assay

Cells were seeded to obtain over 8 x 10^6^ cells, treated and labelled with BrdU (B5002; Sigma) for 1h before harvesting them. After 3 PBS washes, cells were trypsinised and collected in cold PBS, washed twice with PBS, and fixed with ethanol 75% (final concentration) at least 2h at −20°C. Cells could be stored at −20°C for up to one week. Cells were counted and stained with PI/RNAse (550825; BD Pharmigen) following the manufacturer’s recommendations. Cells were then counted in order to obtain at least 20.000 cells for each S phase fraction. Cells were sorted into 4 S-phase fractions (S1-S4) using FACS Aria II (Cytometry and Cell Sorting Core Facility, IDIBAPS). Sorted cells were kept on ice after sorting. After centrifugation, cells were washed once with PBS and lysed with SDS-PK buffer (50 mM Tris-HCl; 10 mM EDTA; 1 M NaCl; 0.5% SDS) with 0.2 mg/mL proteinase K (A3830,0100, PanReac Applichem) and 0.05 mg/mL glycogen for 2 hours at 56°C, at the ratio of 1 mL per 100,000 cells. The samples were aliquoted into 200 uL volumes (equivalent to 20,000 cells each). Lysates could be stored at −20°C for several months. DNA from each aliquot was purified using Zymo Quick DNA Microprep Kit (Zymo Research, D3020) according to the manufacturer’s instructions. Purified DNA was fragmented using a Bioruptor standard UCD-200 (DIAGENODE) to obtain fragments from 500 to 1000 bp. DNA fragments were then denatured at 95°C for 5 minutes and immediately chilled on ice for 2 minutes. BrdU incorporated was immunoprecipitated with Dynabeads protein G (100003D, ThermoFisher) following the manufacturer’s recommendations using anti-BrdU (347580, BD Biosciences) was incubated for 30 minutes on a rotating orbital.DNA recovered from dynabeads was purified with a DNA clean and Concentrator-5 kit (Zymo Research, D4013). Amplification of the recovered DNA was carried out using a whole genome amplification kit (WGA2, Sigma-Aldrich) according to the manufacturer’s recommendations. The final purification of amplified DNA was performed with the DNA clean and Concentrator-5 kit (Zymo Research, D4013) and quantified using NanoDrop one (260nm). DNA was then analysed by qPCR.

### Retrotranscription

RNA extraction was performed using RNeasy Mini Kit (74104, QUIAGEN) following the manufacturer’s recommendations, eluting in the final step with RNAse free water. The following retrotranscription was done with commercial kit (4368814, ThermoFischer). The cDNA obtained was analysed with qPCR.

### qPCR

For replication timing analysis, qPCR was conducted in 384-well plates with a final reaction volume of 5 µL containing 1 ng of DNA per well. The qPCR was run on a QS5 system (ThermoFischer) with the following program: 50°C for 2 min, 95°C for 10 min, followed by 50 cycles (95°C for 15 seconds, 60°C for 30 seconds, 72°C for 30 seconds) and a melt curve (95°C for 15 seconds, 55°C for 1 min, with a ramp of 0.075°C/sec up to 95°C for 15 seconds). SYBR green detection was normalized to mitochondrial DNA for each S phase fraction. The relative detection for each stage of the S phase was calculated as a proportion of each stage respectively of the total detection. Replication timing shift was calculated as *RTshift_Gene_*= *RT_Gene_*_, *Treated*_ − *RT_Gene_*_, *Control*_; considering *RT_Gene_* = ((4 · *S*1 + 3 · *S*2 + 2 · *S*3 + 1 · *S*4) − 250)⁄150.

For cDNA analysis, qPCR was performed on a LightCycler (Roche) in 96-well plates with a final volume of 1.5 µL using a 1/20 dilution of the cDNA. The program used was: 95°C for 10 min, followed by40 cycles (95°C for 30 seconds, 60°C for 15 seconds, 72°C for 30seconds) and a melt curve (95°C for 10 seconds - 65°C for 60 seconds, 97°C for 1 second continuous). *GADPH* served as a loading control.

The primers used are listed in Table 1.

**Table 1.**
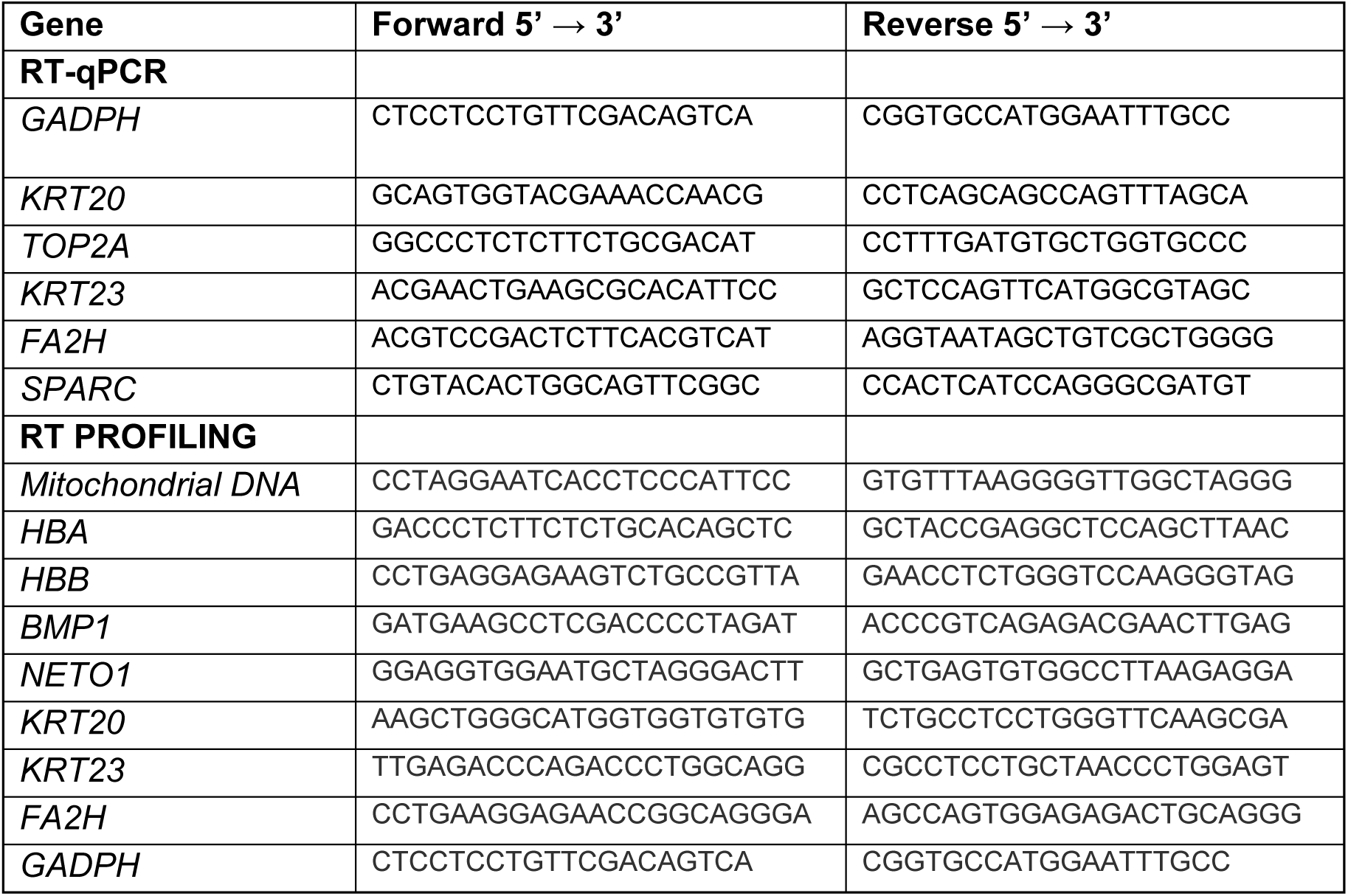
List of primers used in this study.

### Assay for transposase accessible chromatin sequencing (ATAC-seq) assay

ATAC-seq was done according to Buenrostro et al.(Buenrostro *et al*, 2016) until transposition reaction. Once purified, DNA library generation and sequencing steps were performed into Genomic Unit of “Centre de regulació Genòmica”. Libraries were sequenced with NovaSeq 6000 Dx (Ilumina) 2×50M mapped reads. Data were processed following the pipeline of ATACseq from Galaxy project. Briefly, three independent experiments were analysed together to increase statistical power. Reads were trimmed and mapped using bowtie2 to the human reference genome h38, removing reads with quality scores below 30, mitochondrial reads and duplicates. Reads were checked in size, showing a peak around 200 bp. Open regions were identified with MACS2 peak calling using the broad mode, following the ENCODE recommendations. Resulting archives were converted to BigWig and visualized with the UCSC genome browser. Subsequently, the comparison between different experimental groups was performed identifying common and condition-specific peaks. Peaks were considered overlapped if intersected on at least two bp. Quantitative changes in chromatin accessibility were analysed comparing the peaks of individual replicates for each condition. Statistical significance and fold changes in chromatin accessibility were calculated using a t-test. Genomic mapping of the different peaks was done with ChIPseeker in Galaxy platform. Data were analysed with tidyverse (v2.0.0) UpSetR (v1.4.0) packages in R environment (R version 4.4.1).

### Data analysis

The data analysis was made with GraphPad Prism (version 10.3.1 for Windows, GraphPad Software, Boston, Massachusetts USA, www.graphpad.com) or R studio (RStudio 2024.09.0+375) as indicated.

## ABBREVIATIONS

CRC: colorectal cancer
CldU: 5-chloro-2’deoxyuridine
DNA: deoxyribonucleic acid
HU: hydroxyurea
IdU: 5-iodo-2’deoxyuridine
RD: replication domains

## AUTHOR CONTRIBUTIONS

A.P-V and F.U. contributed equally to this work; Conceptualization: A.P-V., F.U. C.M. and N.A.; Methodology: A.P., F.U, S.F. and P.LL-A.; Investigation: A.P-V., P.U. and S.F; Data analysis A.P-V., F.U, and P.LL-A.; Writing—Review & Editing: A.P-V, F.U., C.M. and N.A; Project administration: N.A.; Supervision: C.M. and N.A; Funding acquisition: N.A and C.M. All authors have read and agreed to the published version of the manuscript.

## FUNDING

This research was funded by: Proyecto PID2019-105483RB-I00 financiado por MICIU/AEI /10.13039/501100011033 and Proyecto PID2022-138728OB-I00 financiado por MICIU/AEI/10.13039/501100011033 y por FEDER, UE to N.A., a PREDOCS-UB fellowship co-funded with BancoSantander for A.P-V., a FPU fellowships from Ministerio de Educación, Cultura y Deporte for F.U. and a FI fellowship from Generalitat de Catalunya to S.F. This study was supported by grants to CM from the Ministerio de Ciencia e Innovación and Agencia Estatal de Investigación (PID2020-118768RJ-I00; AEI/10.13039/501100011033), a Ramon y Cajal fellowship (RYC2022-035576-I) and a research grant from the Asociación Española Contra el Cáncer AECC (LABAE222994MAUV).

## ACKNOWLEDGMENTS

We thank the personnel of the Advanced Microscopy Unit of CCiTUB (Campus Clinic) for their help in setting up the image acquisition and analysis.

## SUPPLEMENTARY METHODS

### Cell viability assay

Cells were trypsinised and centrifuged at 660 rcf for 5 minutes at 4°C. Cells were stained with annexin-PI solution (50 μL Annexin buffer 10X; 2 μl APC-AnnexinV 550474 BD; 0.025 μL PI 1 μg/μL, P4864 Sigma) prepared with Annexin buffer 10x (2.5 mM CaCl2; 1.4 M NaCl; 1.4 M Hepes) and incubated for 10 minutes in the dark at 4°C. The samples were processed by flow cytometry within the following hour.

### Flow cytometry

Cells were harvested and fixed in 70% ethanol for at least 2 h at −20°C before immunostaining and flow cytometry. Combined analysis of DNA content with propidium iodide (PI), BrdU (anti-BrdU, Abcam, Cambridge, United Kingdom, ab6326; 1/250) positive population and the MPM2 (anti-MPM2, Millipore, Burlington, MA, USA, #05– 368; 1/250) positive population were performed with a BD FACSCanto II (Cytometry and Cell Sorting Core Facility, IDIBAPS) as previously described (Ercilla *et al*, 2016).

## SUPPLEMENTARY FIGURES

**Sup.Fig1. Inhibition of CDK2 slightly sensitizes HCT116 cells to HU-induced replication stress and delays S phase fulfilment.**

**A.** Experimental design of (B). HU, hydroxyurea; CDK2i, CDK2 inhibitor type II.

**B.** Cell viability analysed by Annexin V in HCT116 cells upon HU-induced replication stress treated or not with CDK2 inhibitor (CDK2i) during recovery with reduces the viability of the cells (N=3; left) even detected after 8 hours of recovery in fresh medium (N=1; right) studied by Annexin V.

**C.** Experimental design of (D) and (E). HU, hydroxyurea; CDK2i, CDK2 inhibitor type II.

**D.** Cells were treated as in A, and before harvesting they were incubated for 20 min with BrdU to detect cells in S phase.

**Sup.Fig.2. HCT116 cells enter mitosis 11 hours following HU-induced replication stress.**

**A.** Experimental design. BrdU, 5-bromo-2’deoxyuridine; HU, hydroxyurea.

**B.** Mitotic cells detected with MPM2 labelling by flow cytometry. BrdU positive cells are those that were in S phase when HU was added and consequently have been in S phase during the 14h of HU treatment.

**Sup.Fig.3. Nuclear area is increased with HU treatment and the increase maintained during the following cell cycles.**

**A-C**: Analysis of the nuclear area. Replicated experiments of Fig4. Data are presented as a boxplot (mean plus second and third quartile as a box) with a violin plot to assess de cell area distribution. Statistical significances were calculated with a Kruskal-Wallis’ test (non-parametric ANOVA) followed by a Dunn test for multiple comparison’s post hoc test. Ns, not-significative; *p < 0.05; **p < 0.01; ***p < 0.001; ****p < 0.0001; compared to control conditions.

**Sup.Fig.4. UCSC genome browser viewer of selected genes.** The first track highlights the GENCODE Genes version 47, September 2024. From the 2nd to the 7th tracks, the comparisons obtained in our data between the different conditions are shown (see Fig 4.). The 8th track shows the regulatory elements and gene interaction comprised in GenHancer embedded in GeneCards and present the regulatory elements (GeneHancers), gene transcription start sites (TSS), interactions or associations between regulatory elements and genes, and clustered interactions by gene target or GeneHancer. The highlighted area represents the peak detected related to each gene. Note that it could be more than one peak. **A.** *KRT20* and *KRT23*. **B.** *NETO1*. **C.** *TOP2A*. **D.** SPARC**. E.** BMP1. **F.** FA2H. **G.** GADPH.

**Sup.Fig.5. Replication stress induced by HU alters the transcription of different genes.**

**A.** Experimental design. Hydroxyurea (HU; 10mM). CDK2 type II inhibitor (CDK2i; 10mM).

**B.** qPCR results presented as a fold change regarding control condition. Data are presented as columns with mean ± SEM. Significance was calculated as a Kruskal-Wallis’ test (non-parametric ANOVA) followed by a Dunn test for multiple comparison’s post hoc test. *p<0.05; **p<0.01; ***p<0.005; ****p<0.001.

## Notes

### Competing Interest Statement

The authors have declared no competing interest.

## REFERENCES

Aird KM, Zhang G, Li H, Tu Z, Bitler BG, Garipov A, Wu H, Wei Z, Wagner SN, Herlyn M, et al (2013) Suppression of Nucleotide Metabolism Underlies the Establishment and Maintenance of Oncogene-Induced Senescence. Cell Rep 3: 1252–1265

Akerman I, Kasaai B, Bazarova A, Sang PB, Peiffer I, Artufel M, Derelle R, Smith G, Rodriguez-Martinez M, Romano M, et al (2020) A predictable conserved DNA base composition signature defines human core DNA replication origins. Nat Commun 11: 1–15

Aladjem MI & Redon CE (2017) Order from clutter: Selective interactions at mammalian replication origins. Nat Rev Genet 18: 101–116

Alver RC, Chadha GS & Blow JJ (2014) The contribution of dormant origins to genome stability: From cell biology to human genetics. DNA Repair (Amst) 19: 182–189

Bester AC, Roniger M, Oren YS, Im MM, Sarni D, Chaoat M, Bensimon A, Zamir G, Shewach DS & Kerem B (2011) Nucleotide Deficiency Promotes Genomic Instability in Early Stages of Cancer Development. Cell 145: 435–446

Boyer A-S, Walter D & Sørensen CS (2016) DNA replication and cancer: From dysfunctional replication origin activities to therapeutic opportunities. Semin Cancer Biol 37–38: 16–25

Branzei D & Foiani M (2009) The checkpoint response to replication stress. DNA Repair (Amst) 8: 1038–46

Branzei D & Foiani M (2010) Maintaining genome stability at the replication fork. Nat Rev Mol Cell Biol 11: 208–19

Buenrostro J, Wu B, Chang H & Greenleaf W (2016) ATAC-seq method. Curr Protoc Mol Biol 2015: 1–10

Burrell RA, McClelland SE, Endesfelder D, Groth P, Weller M-C, Shaikh N, Domingo E, Kanu N, Dewhurst SM, Gronroos E, et al (2013a) Replication stress links structural and numerical cancer chromosomal instability. Nature 494: 492–6

Burrell RA, McGranahan N, Bartek J & Swanton C (2013b) The causes and consequences of genetic heterogeneity in cancer evolution. Nature 501: 338–345 doi:10.1038/nature12625 [PREPRINT]

Cayrou C, Ballester B, Peiffer I, Fenouil R, Coulombe P, Andrau JC, Van Helden J & Méchali M (2015) The chromatin environment shapes DNA replication origin organization and defines origin classes. Genome Res 25: 1873–1885

Chagin VO, Casas-Delucchi CS, Reinhart M, Schermelleh L, Markaki Y, Maiser A, Bolius JJ, Bensimon A, Fillies M, Domaing P, et al (2016) 4D Visualization of replication foci in mammalian cells corresponding to individual replicons. Nat Commun 7: 11231

Courtot L, Bournique E, Maric C, Guitton-Sert L, Madrid-Mencía M, Pancaldi V, Cadoret JC, Hoffmann JS & Bergoglio V (2021) Low replicative stress triggers cell-type specific inheritable advanced replication timing. Int J Mol Sci 22

Donley N & Thayer MJ (2013) DNA replication timing, genome stability and cancer. Late and/or delayed DNA replication timing is associated with increased genomic instability. Semin Cancer Biol 23: 80–89

Dunham I, Kundaje A, Aldred SF, Collins PJ, Davis CA, Doyle F, Epstein CB, Frietze S, Harrow J, Kaul R, et al (2012) An integrated encyclopedia of DNA elements in the human genome. Nature 489: 57–74

Ercilla A, Llopis A, Feu S, Aranda S, Ernfors P, Freire R & Agell N (2016) New origin firing is inhibited by APC/CCdh1activation in S-phase after severe replication stress. Nucleic Acids Res 44: 4745–4762

Errico A & Costanzo V (2010) Differences in the DNA replication of unicellular eukaryotes and metazoans: Known unknowns. EMBO Rep 11: 270–278

Feu S, Unzueta F, Ercilla A, Perez-Venteo A, Jaumot M & Agell N (2022a) RAD51 is a druggable target that sustains replication fork progression upon DNA replication stress. PLoS One 17: 1–20

Feu S, Unzueta F, Ercilla A, Perez-Venteo A, Jaumot M & Agell N (2022b) RAD51 is a druggable target that sustains replication fork progression upon DNA replication stress. PLoS One 17: 1–20

Foti R, Gnan S, Cornacchia D, Dileep V, Bulut-Karslioglu A, Diehl S, Buness A, Klein FA, Huber W, Johnstone E, et al (2016) Nuclear Architecture Organized by Rif1 Underpins the Replication-Timing Program. Mol Cell 61: 260–273

Fragkos M, Ganier O, Coulombe P & Méchali M (2015) DNA replication origin activation in space and time. Nat Rev Mol Cell Biol 16: 360–74

Gaillard H, García-Muse T & Aguilera A (2015) Replication stress and cancer. Nat Rev Cancer 15: 276–289

Ge XQ, Jackson DA & Blow JJ (2007) Dormant origins licensed by excess Mcm2-7 are required for human cells to survive replicative stress. Genes Dev 21: 3331–41

Hanahan D (2022) Hallmarks of Cancer: New Dimensions. Cancer Discov 12: 31–46 doi:10.1158/2159-8290.CD-21-1059 [PREPRINT]

Hanahan D & Weinberg RA (2011) Hallmarks of Cancer: The Next Generation. Cell 144: 646–674

Hiratani I, Ryba T, Itoh M, Rathjen J, Kulik M, Papp B, Fussner E, Bazett-Jones DP, Plath K, Dalton S, et al (2010) Genome-wide dynamics of replication timing revealed by in vitro models of mouse embryogenesis. Genome Res 20: 155–169

Hiratani I, Ryba T, Itoh M, Yokochi T, Schwaiger M, Chang CW, Lyou Y, Townes TM, Schübeler D & Gilbert DM (2008) Global reorganization of replication domains during embryonic stem cell differentiation. PLoS Biol 6: 2220–2236

Hsu CL, Chong SY, Lin CY & Kao CF (2021) Histone dynamics during DNA replication stress. J Biomed Sci 28 doi:10.1186/s12929-021-00743-5 [PREPRINT]

Jasencakova Z, Scharf AND, Ask K, Corpet A, Imhof A, Almouzni G & Groth A (2010) Replication Stress Interferes with Histone Recycling and Predeposition Marking of New Histones. Mol Cell 37: 736–743

Jodkowska K, Pancaldi V, Rigau M, Almeida R, Fernández-Justel JM, Graña-Castro O, Rodríguez-Acebes S, Rubio-Camarillo M, Carrillo-De Santa Pau E, Pisano D, et al (2022) 3D chromatin connectivity underlies replication origin efficiency in mouse embryonic stem cells. Nucleic Acids Res 50: 12149–12165

Kang S, Kang MS, Ryu E & Myung K (2018) Eukaryotic DNA replication: Orchestrated action of multi-subunit protein complexes. Mutation Research - Fundamental and Molecular Mechanisms of Mutagenesis 809: 58–69

Lande-Diner L, Zhang J & Cedar H (2009) Shifts in Replication Timing Actively Affect Histone Acetylation during Nucleosome Reassembly. Mol Cell 34: 767–774

Macheret M & Halazonetis TD (2015) DNA replication stress as a hallmark of cancer. Annu Rev Pathol 10: 425–48

Muñoz S & Méndez J (2016) DNA replication stress: from molecular mechanisms to human disease. Chromosoma 126: 1–15

Nakatani T, Schauer T, Altamirano-Pacheco L, Klein KN, Ettinger A, Pal M, Gilbert DM & Torres-Padilla ME (2024) Emergence of replication timing during early mammalian development. Nature 625: 401–409

Neelsen KJ, Zanini IMY, Mijic S, Herrador R, Zellweger R, Chaudhuri AR, Creavin KD, Julian Blow J & Lopes M (2013) Deregulated origin licensing leads to chromosomal breaks by rereplication of a gapped DNA template. Genes Dev 27: 2537–2542

Nieto-Soler M, Morgado-Palacin I, Lafarga V, Lecona E, Murga M, Callen E, Azorin D, Alonso J, Lopez-Contreras AJ, Nussenzweig A, et al (2016) Efficacy of ATR inhibitors as single agents in Ewing sarcoma. Oncotarget 7: 58759–58767

Pope BD, Ryba T, Dileep V, Yue F, Wu W, Denas O, Vera DL, Wang Y, Hansen RS, Canfield TK, et al (2014) Topologically associating domains are stable units of replication-timing regulation. Nature 515: 402–405

Primo LMF & Teixeira LK (2020) Dna replication stress: Oncogenes in the spotlight. Genet Mol Biol 43: 1–14

Rivera-Mulia JC & Gilbert DM (2016a) Replication timing and transcriptional control: Beyond cause and effect - part III. Curr Opin Cell Biol 40: 168–178

Rivera-Mulia JC & Gilbert DM (2016b) Replicating Large Genomes: Divide and Conquer. Mol Cell 62: 756–765 doi:10.1016/j.molcel.2016.05.007 [PREPRINT]

Ryba T, Hiratani I, Lu J, Itoh M, Kulik M, Zhang J, Schulz TC, Robins AJ, Dalton S & Gilbert DM (2010) Evolutionarily conserved replication timing profiles predict long-range chromatin interactions and distinguish closely related cell types. Genome Res 20: 761–770

Sequeira-Mendes J, Díaz-Uriarte R, Apedaile A, Huntley D, Brockdorff N & Gómez M (2009) Transcription initiation activity sets replication origin efficiency in mammalian cells. PLoS Genet 5

Smith OK, Kim R, Fu H, Martin MM, Lin CM, Utani K, Zhang Y, Marks AB, Lalande M, Chamberlain S, et al (2016) Distinct epigenetic features of differentiation-regulated replication origins. Epigenetics Chromatin 9

Sugimoto N, Maehara K, Yoshida K, Ohkawa Y & Fujita M (2018) Genome-wide analysis of the spatiotemporal regulation of firing and dormant replication origins in human cells. Nucleic Acids Res 46: 6683–6696

Técher H, Koundrioukoff S, Nicolas A & Debatisse M (2017) The impact of replication stress on replication dynamics and DNA damage in vertebrate cells. Nat Rev Genet 18: 535–550

Ubhi T & Brown GW (2019) Exploiting DNA Replication Stress for Cancer Treatment. Cancer Res 78: 1–11

Wilson KA, Elefanty AG, Stanley EG & Gilbert DM (2016) Spatio-temporal re-organization of replication foci accompanies replication domain consolidation during human pluripotent stem cell lineage specification. Cell Cycle 15: 2464–2475

Yamazaki S, Ishii A, Kanoh Y, Oda M, Nishito Y & Masai H (2012) Rif1 regulates the replication timing domains on the human genome. EMBO Journal 31: 3667–3677

Zhao PA, Rivera-Mulia J-C & Gilbert DM (2017) Replication Domains: Genome Compartmentalization into Functional Replication Units Masai H & Foiani M (eds) Springer Nature

