## Supplementary figures and images for "Adaptive replication origin activation alters chromatin dynamics and stability in cancer cells"

### Supplementary Figures

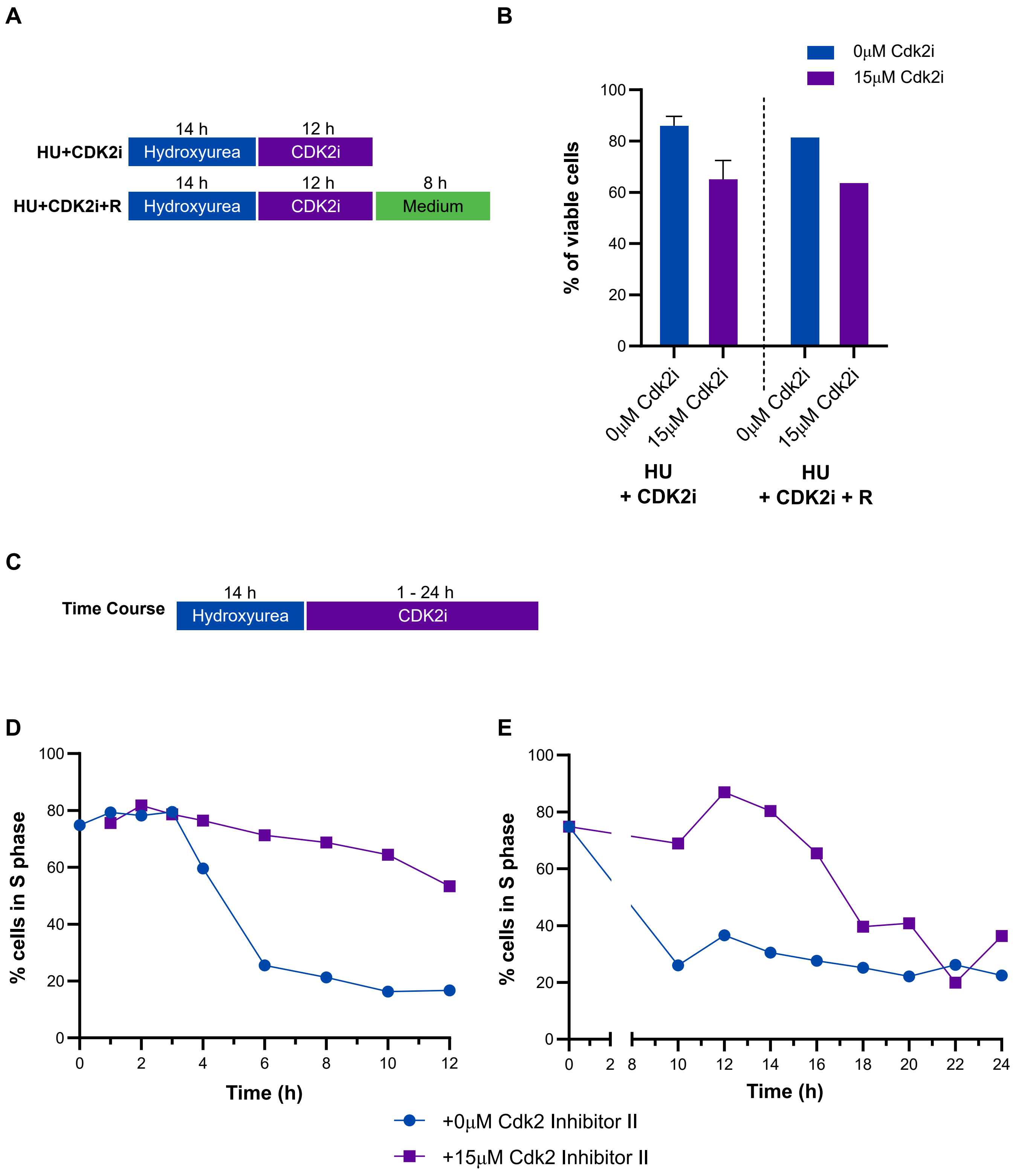

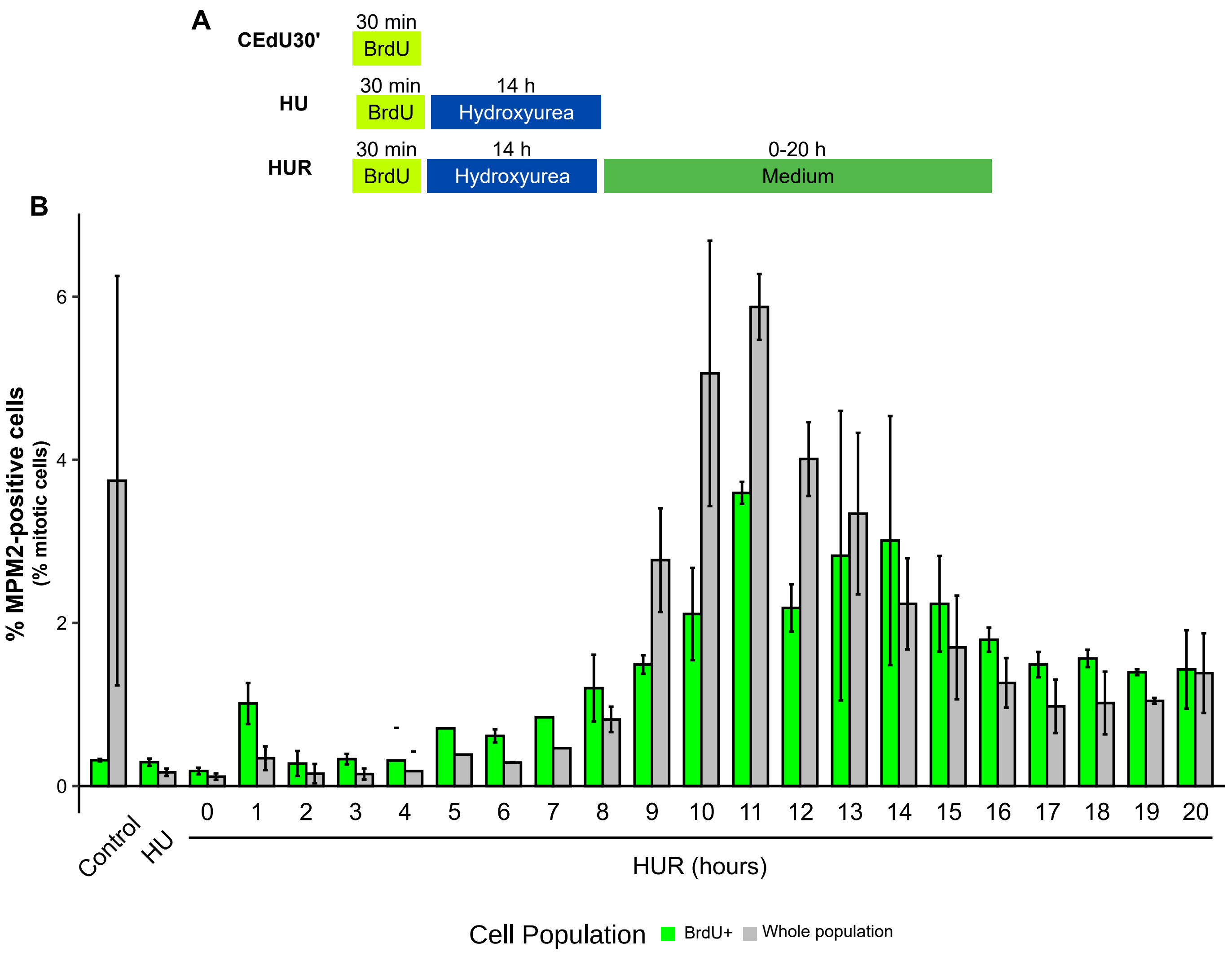

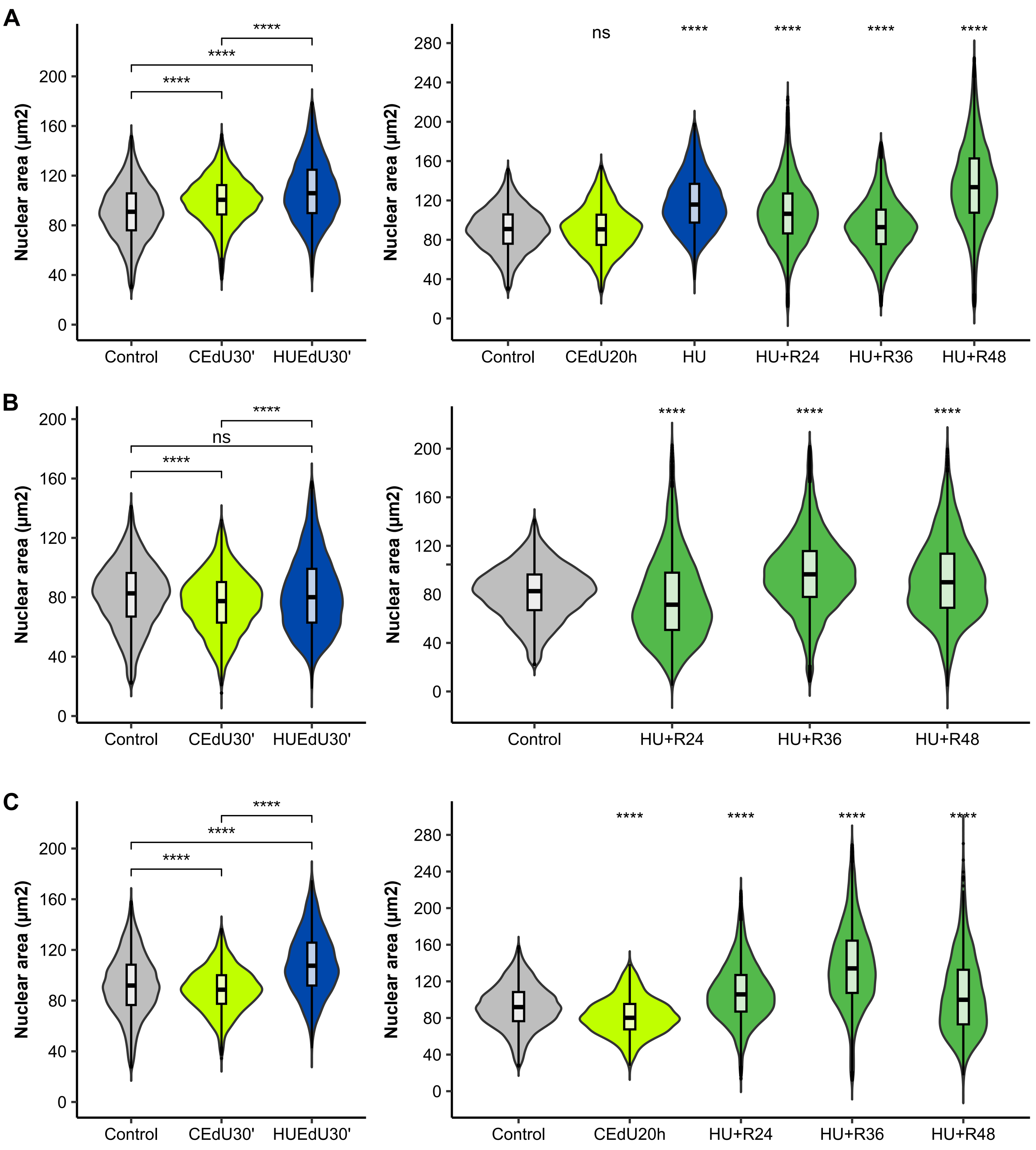

SUP. FIGURE 4

A

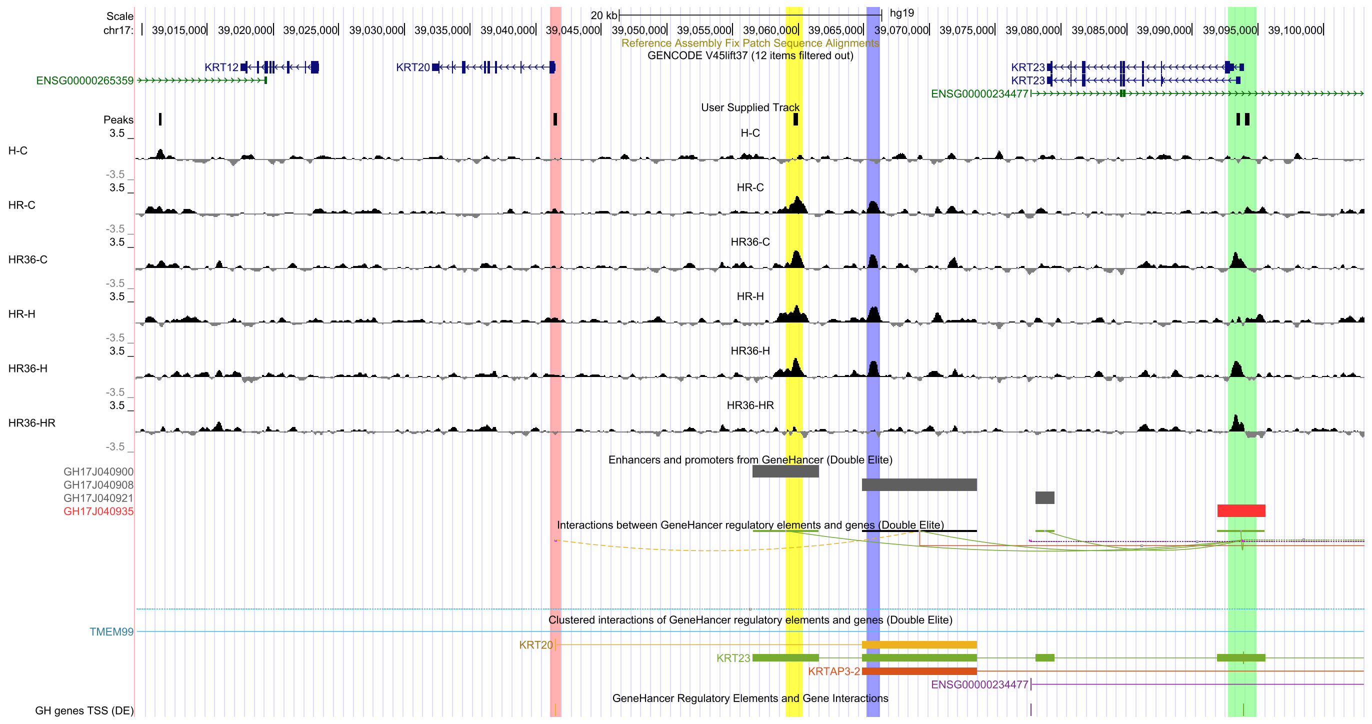


B

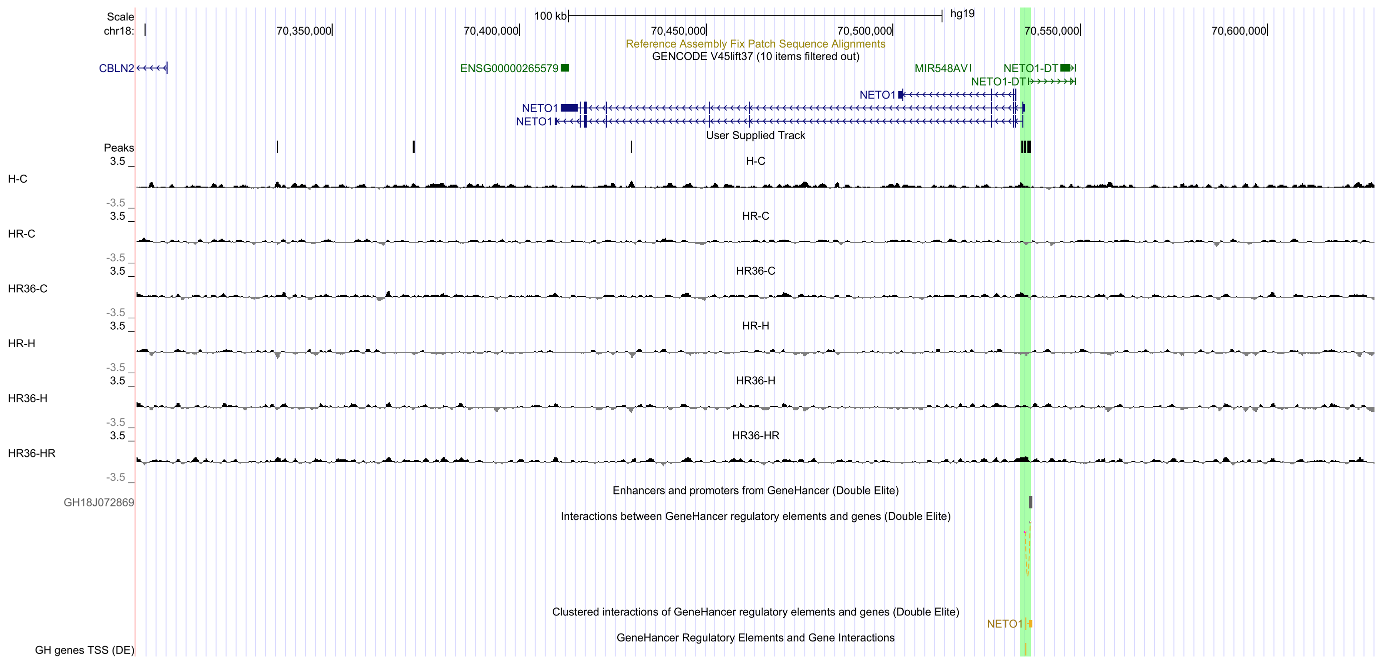

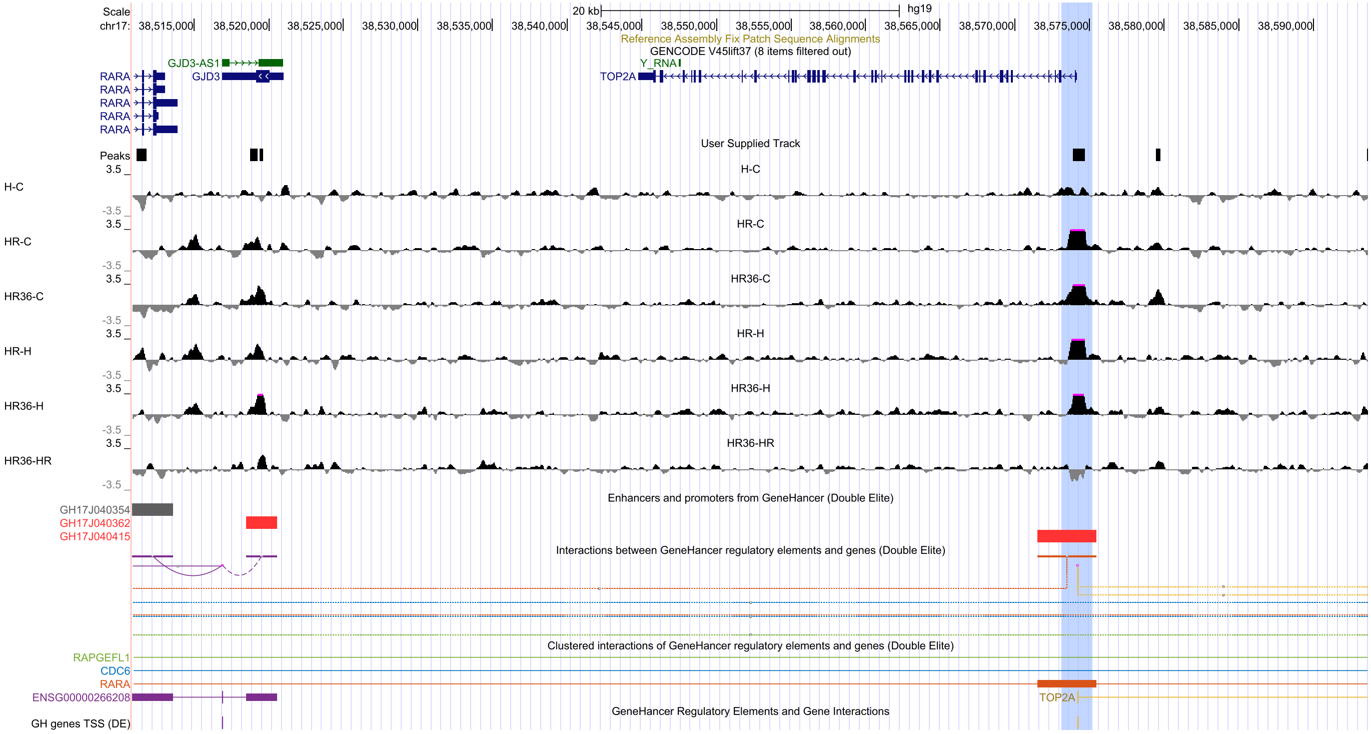

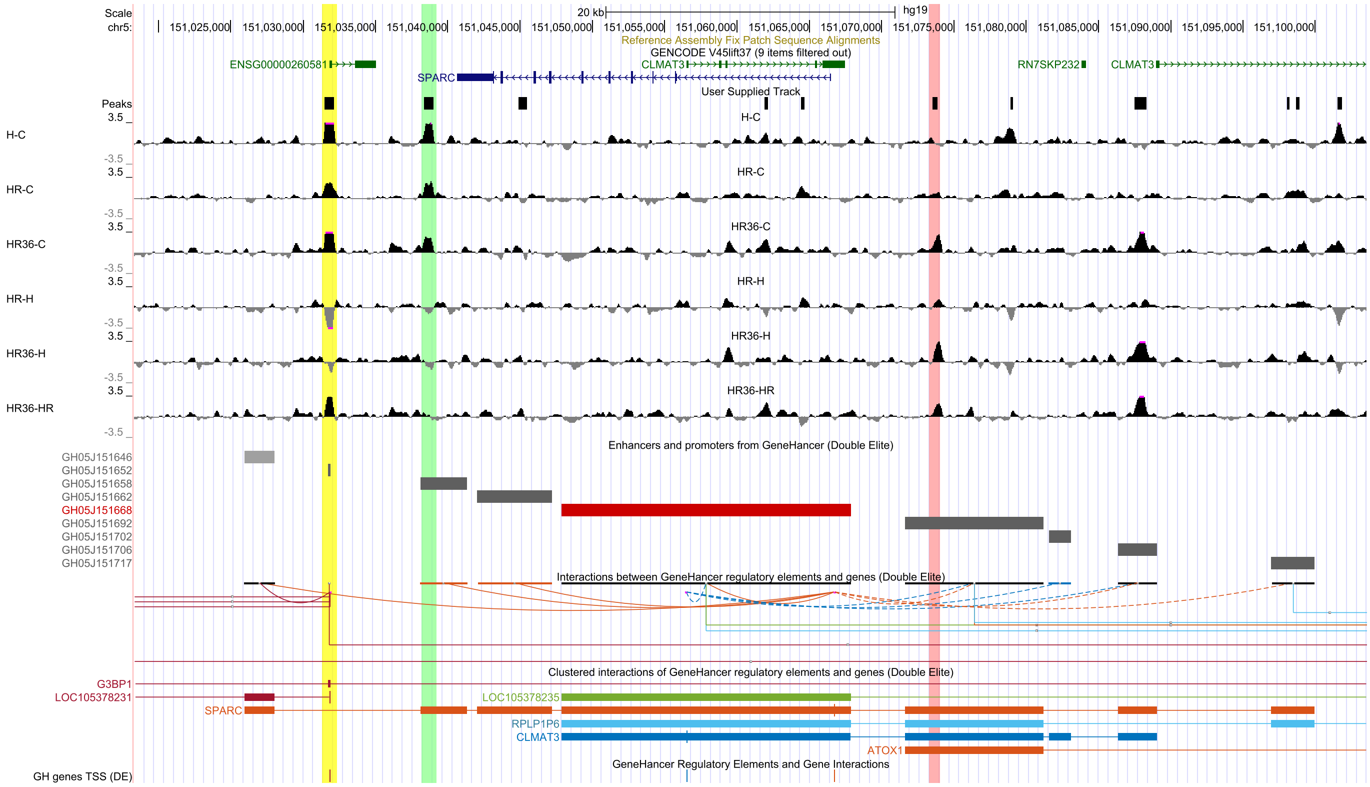

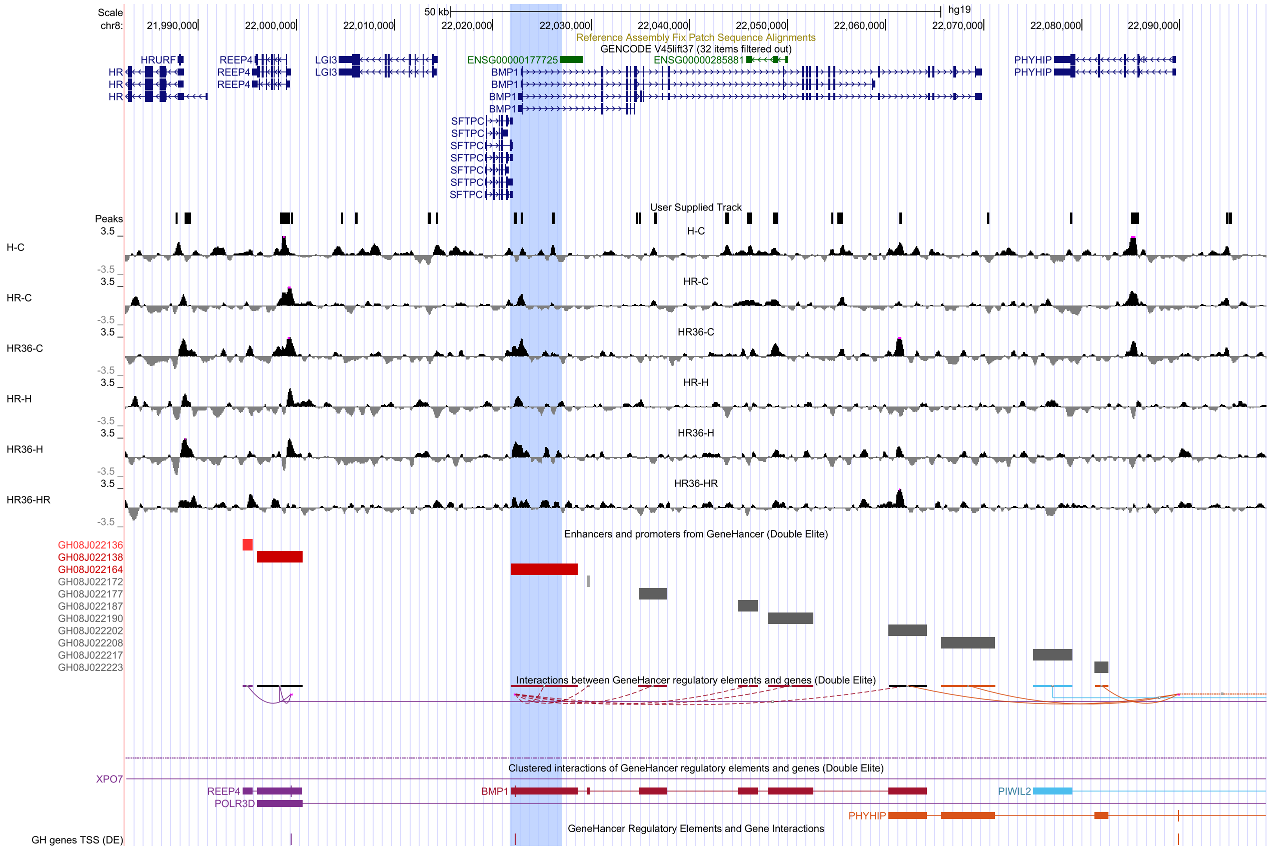

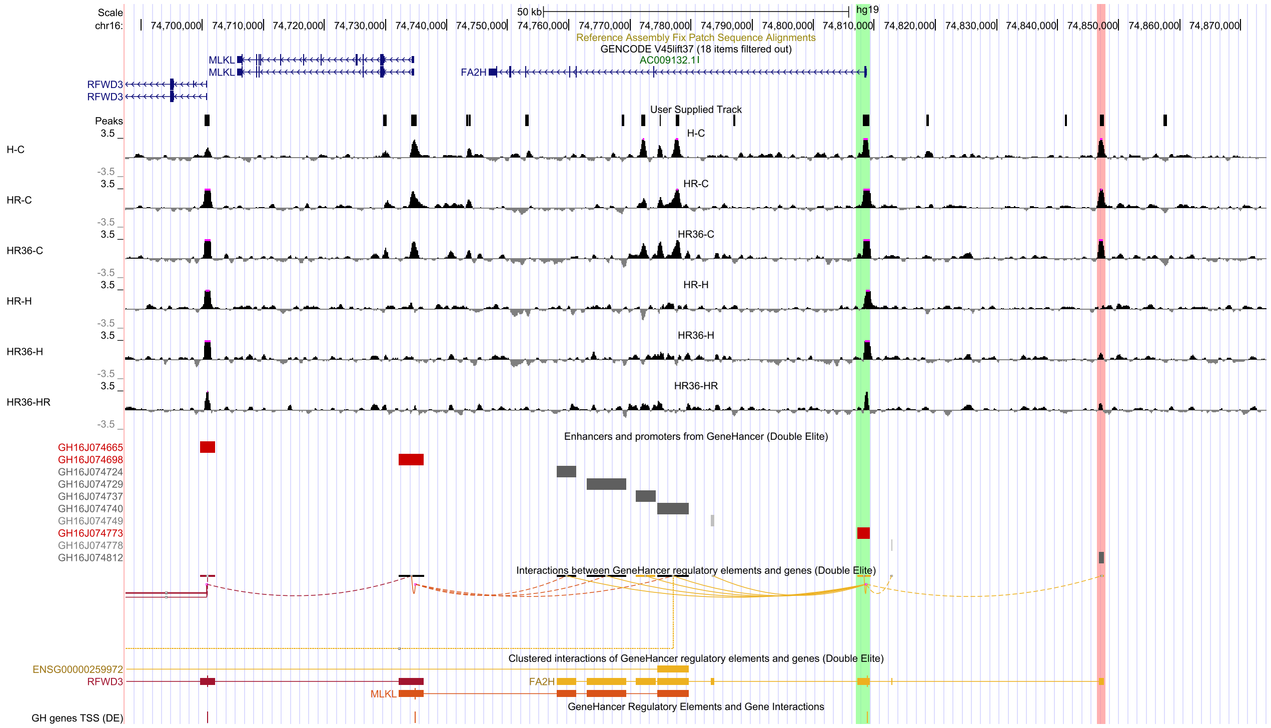

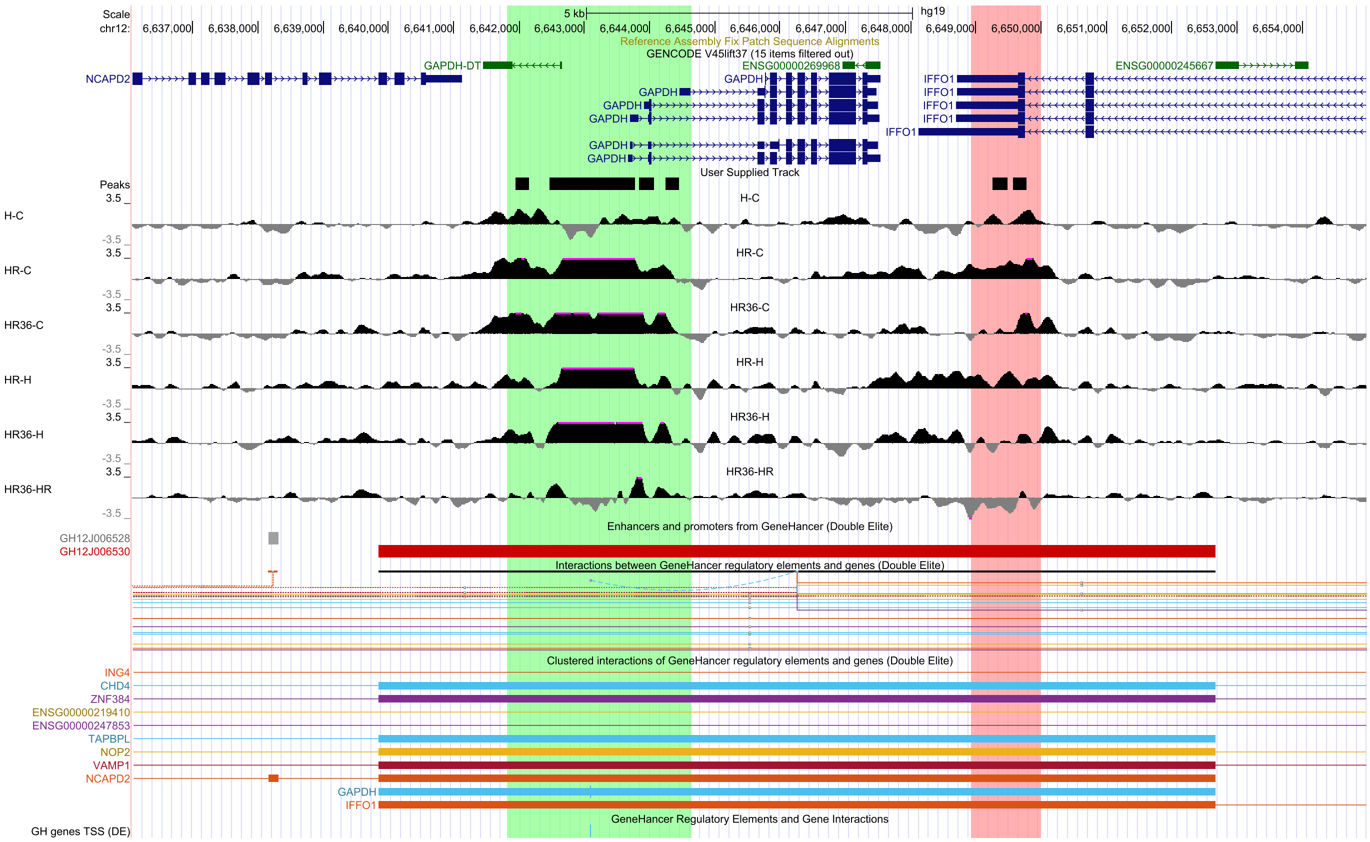

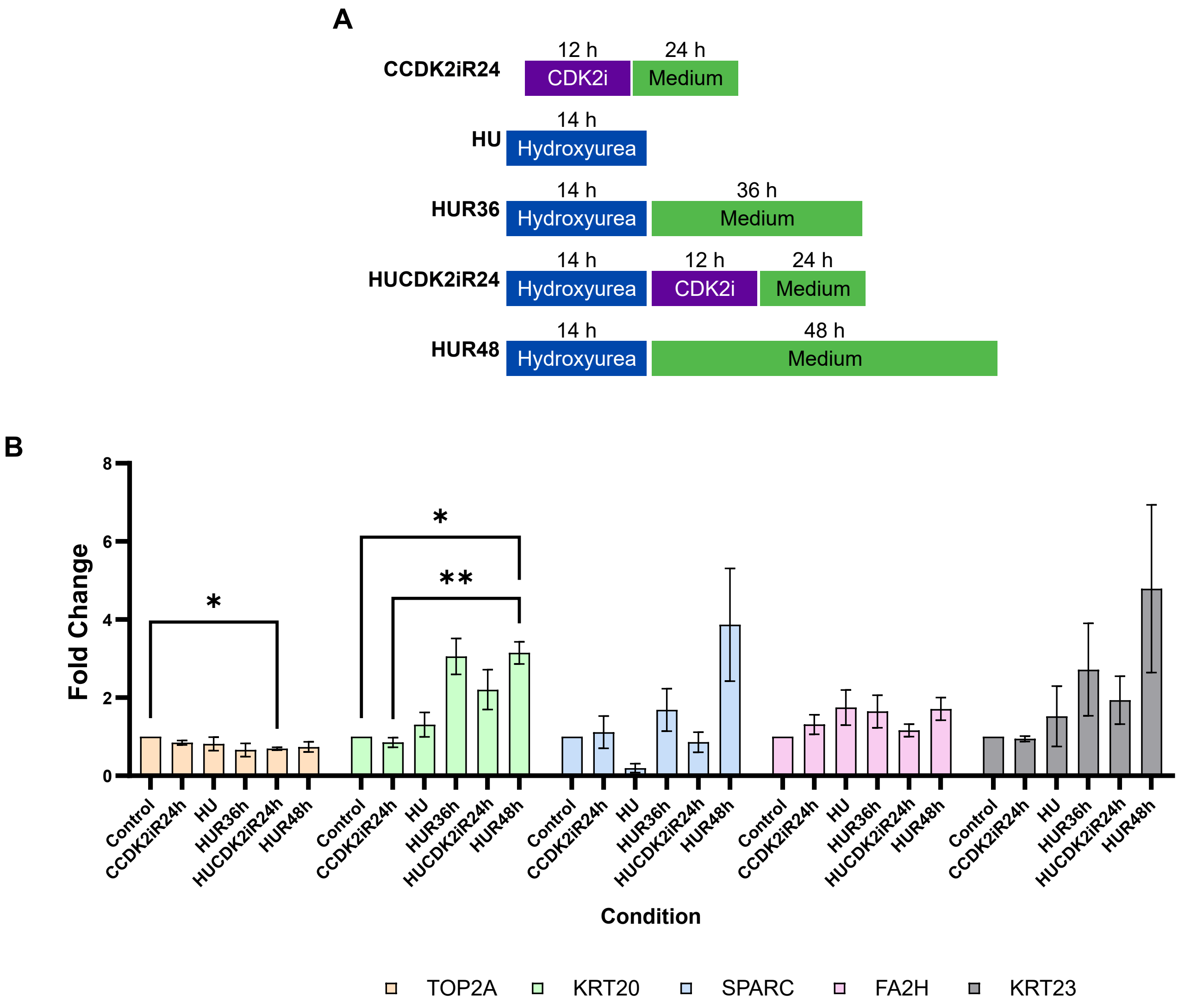
